# Ecological drivers of body size evolution and sexual size dimorphism in short-horned grasshoppers (Orthoptera: Acrididae)

**DOI:** 10.1101/119560

**Authors:** Vicente García-Navas, Víctor Noguerales, Pedro J. Cordero, Joaquín Ortego

## Abstract

Sexual size dimorphism (SSD) is widespread and variable in nature. Although female-biased SSD predominates among insects, the proximate ecological and evolutionary factors promoting this phenomenon remain largely unstudied. Here, we employ modern phylogenetic comparative methods on 8 subfamilies of Iberian grasshoppers (85 species) to examine the validity of different models of evolution of body size and SSD and explore how they are shaped by a suite of ecological variables (habitat specialization, substrate use, altitude) and/or constrained by different evolutionary pressures (female fecundity, strength of sexual selection, length of the breeding season). Body size disparity primarily accumulated late in the history of the group and did not follow a Brownian motion pattern, indicating the existence of directional evolution for this trait. We found support for the converse of Rensch’s rule across all taxa but not within the two most speciose subfamilies (Gomphocerinae and Oedipodinae), which showed an isometric pattern. Our results do not provide support for the fecundity or sexual selection hypotheses and we did not find evidence for significant effects of habitat use. Contrary to that expected, we found that species with narrower reproductive window are less dimorphic in size than those that exhibit a longer breeding cycle, suggesting that male protandry cannot solely account for the evolution of female-biased SSD in Orthoptera. Our study highlights the need to consider alternatives to the classical evolutionary hypotheses when trying to explain why in certain insect groups males remain small.

---

Body size is a key trait in any organism, as it influences fitness through its effects on reproduction and survival (Fairbairn et al. 2007). Body size can respond to different evolutionary forces in a sex-specific manner and, as a result, this trait often differs between males and females in many taxa (Darwin 1871). Although male-biased sexual size dimorphism (SSD) is the common rule among endotherms (mammals and birds), female-biased SSD predominates among insects (e.g. Elgar 1991, Cheng and Kuntner 2014, Hochkirch and Gröning 2008, Bidau et al. 2016). In those species in which females are larger than males, it is assumed that natural selection on female body size (*via* increased fecundity) overrides sexual selection (through competition advantages during mate acquisition) on male body size. However, many other ecological pressures (e.g. habitat, substrate use, length of life cycle) can determine body size evolution in one or both sexes and contribute to shape observed patterns of SSD (Fairbairn et al. 2007, Blanckenhorn et al. 2007a, Fairbairn 2013).

Body size variation among related species often follows evolutionary patterns that are remarkably consistent across taxa. According to the Rensch’s rule, SSD increases with body size when males are the larger sex and decreases with size when females are larger (Rensch 1950, Abouheif and Fairbairn 1997, Fairbairn 1997). In contrast, if selection pressures on females are the main driver of SSD evolution, then SSD should increase with average body size in female-biased SSD species (the converse of Rensch’s rule). Alternatively, SSD can remain isometric when body size changes in males and females at the same rate, a plausible scenario when multiple evolutionary forces (e.g. sexual selection and fecundity selection) act synergically with no overall general trend. Another broad-scale pattern of body size variation is the Bergmann’s rule, which posits the existence of a positive relationship between body size and latitude and altitude, with smaller individuals being found at lower latitudes/altitudes where temperatures are generally higher (Mayr 1956). However, ectotherms often follow the converse of Bergmann’s rule with larger individuals and species at lower latitudes and altitudes (reviewed in Shelomi 2012). The most likely explanation for this inverted cline is the interaction between the length of the growing season (which decreases as latitude and altitude increase) and the time available to complete development. Slowly developing insects may compensate for a short season by decreasing development time, that is, by reaching the adult stage at a smaller size (e.g. reducing the number of larval instars; Blanckenhorn and Demont 2004). However, some species do not exhibit protandry and both sexes reach adulthood at about the same time but at different sizes (Blanckenhorn et al. 2007b).

In Orthoptera, females are usually larger than males. Hochkirch and Gröning (2008) reported that virtually all of the 1106 Caelifera species analyzed showed female-biased SSD (see also Bidau et al. 2016 for a review). From the female perspective, it seems to be clear that a large body size may confer an advantage in terms of increased fecundity (e.g. Cueva del Castillo and Núñez-Farfán 1999; Whitman 2008). Conversely, small males may benefit from early maturity, greater agility, or lower predation rates (see Whitman 2008 and references therein; see also Blanckenhorn 2000). When the evolution of male and female body size follows divergent evolutionary trajectories, it leads to a decoupling of male and female size evolution.

However, absolute decoupling is rather unlikely because genetic correlations between males and females will tend to constrain independent size evolution of both sexes. Body size decoupling has been suggested as the main cause for the existence of extremely female-biased SSD in spiders, a taxonomic group that has dominated studies on arthropods in this respect (Hormiga et al. 2000, Kuntner and Coddington 2009, Kuntner and Elgar 2014). On the contrary, there is a paucity of interspecific studies on SSD in Orthopterans even though they are fairly abundant, easy to collect, and have large geographic distributions, which makes them an ideal model system to address these questions.

In this study, we employ phylogenetic comparative methods to examine the evolution of body size in short-horned grasshoppers (Orthoptera: Caelifera: Acrididae) and test how this trait co-varies with SSD through evolutionary time. To this end, we constructed a phylogeny comprising a representative sample (*n* = 85 taxa) of all extant species present in the Iberian Peninsula (Presa et al. 2007), including slant-faced grasshoppers (subfamily Gomphocerinae, 48 spp.), band-winged grasshoppers (Oedipodinae, 19 spp.), spur-throated grasshoppers (Catantopinae, 6 spp.) and other minoritary groups (e.g. silent slant-faced grasshoppers, Acridinae). Specifically, we first assessed the tempo and mode of evolution of body size and SSD, which allowed us to infer whether neutral or selective forces drove the evolution of these traits. Second, we examined patterns of body size evolution, including altitudinal clines of body size (Bergmann’s rule) and allometric scalings of male and female body size (Rensch’s rule). Finally, we analyzed the proximate ecological factors (habitat specialization, substrate type, altitude) and evolutionary constraints (female fecundity, strength of sexual selection, length of the breeding season) that may underlie the evolution of male and female body size in these large-bodied insects. Our study constitutes the first to provide a comprehensive view about the factors promoting body size evolution and size dimorphism at the interspecific level in Orthopterans.

## Material and Methods

### Sampling

Grasshoppers were collected during several field campaigns carried out throughout the Iberian Peninsula (see e.g. Ortego et al. 2010, 2012, 2015). Specimens were identified using current identification keys for Palearctic gomphocerine species (Harz 1975, Pascual 1978, Pardo et al. 1991, Lluciá-Pomares 2002), and preserved in 96% ethanol. Our sample (*n* = 85) accounted for three quarters of all extant Acrididae species present in the Iberian Peninsula (83% and 66% of all Gomphocerinae and Oedipodinae species, respectively; Presa et al. 2007). Thus, our sample is representative of the natural variation in this region (580 000 km^2^), including eight of the nine families into which Iberian short-horned grasshoppers have been grouped (Presa et al. 2007). More than half of the species (56%) included in this study are endemic to the Iberian Peninsula or have a distribution restricted to Iberia, France and North Africa.

### Molecular data

Genomic DNA was extracted from the hind femur of the specimens using a salt-extraction protocol (Aljanabi and Martínez 1997). Four mitochondrial gene fragments -(1) cytochrome c oxidase subunit 1 (COI), (2) NADH dehydrogenase subunit 5 (ND5), (3) 12S rRNA (12S) and (4) a fragment containing parts of 16S rRNA (16S)- were amplified by polymerase chain reaction and sequenced. Two nuclear genes were also tested (elongation factor 1 α EF-1 and 28S ribosome unit), but these were discarded because their analysis yielded uninformative topologies with poor resolution (see also Song et al. 2015). For some taxa we failed to obtain reliable sequences, so we complemented our data set with additional sequences retrieved from GenBank. We mainly relied on sequences from two previously published phylogenies: Contreras and Chapco (2006) and Nattier et al. (2011).

### Phylogenetic analyses

Sequences were aligned in MAFFT online version 7 (http://mafft.cbrc.jp/alignment/server/; Katoh and Standley 2013) using a L-INS-i strategy. The alignments of the ribosomal genes (12S, 16S) contained highly unequal distribution of indels and thus, were edited by hand in order to eliminate divergent regions of the alignment and poorly aligned positions. Protein-coding genes (COI, ND5) were checked for stop codons and their correct translation to amino acids in Geneious 8.1.7. The sequences of the four genes (12S, 16S, COI and ND5) were trimmed to 380, 469, 568 and 635 base pairs (bp), respectively, to reduce the proportion of missing data. We used Sequencematrix 1.7.8. (Vaidya et al. 2011) to concatenate single alignment fragments, resulting in a concatenated matrix for a total length of 2055 bp. We were not able to obtain reliable sequences from all four markers for some taxa. However, we opted for adding taxa with missing data since this generally increases phylogenetic accuracy (see Hughes and Vogler 2004). The number of sequences per locus obtained was as follows: 79 for COI, 67 for ND5, 80 for 12S, and 78 for 16S. *Pyrgomorpha conica* (Acridoidea) was included as outgroup in all phylogenetic analyses (Chapco and Contreras 2011).

We performed phylogenetic inference and assessed the support of the clades following two methods: maximum likelihood (ML) and Bayesian inference (BI). We calculated the best-fit models of nucleotide substitution for each of the four genes according to the weighted Akaike Information Criterion (AIC) using jModelTest 0.1.1 (Posada 2008). The TIM2+I+Γ substitution model was selected for 12S, GTR+I+ Γ for 16S, TrN+I+ Γ model for ND5 and TPM3uf+I+ Γ was selected for COI. Maximum likelihood analyses were conducted using GARLI version 2.0 (Zwickl 2006) and PHYML (Guindon and Gascuel 2006). A ML bootstrapping procedure was run in GARLI with two search replicates and 1000 bootstrap replicates. The best-fit substitution model for each partition (gene) was assigned by setting the rate matrix, nucleotide state frequencies and proportion of invariable sites. We selected the best (optimal) tree and obtained support for the clades from a majority-rule (50%) consensus tree computed in PAUP* version 4 (Swofford 2002).

Bayesian analyses were conducted using MrBayes 3.2 (Ronquist et al. 2012) applying a nucleotide substitution model specific to each partition (gene): HKY+I+ Γ for 12S and COI, and GTR +I+ Γ for 16S and ND5. We performed two independent runs with four simultaneous Markov chain Monte Carlo (MCMC) chains, running for a total of 10 million generations and sampled every 1000th generation. The first 25% of samples were discarded as burn-in and a consensus tree from the remaining 7501 trees was built using the “sumt” command before visualization in FigTree v.1.3.1 (Rambaut et al. 2009) (see Supporting Information). A second Bayesian analysis was run with BEAST 1.8.0 (Drummond et al. 2012) in order to estimate the relative time of divergence of the studied taxa. Runs were carried out under an uncorrelated lognormal relaxed-clock model and applying a Yule process as tree prior. Two calibration points were used to impose age constraints on some nodes of the tree allowing us to translate the relative divergence times into absolute ones. We employed as a first calibration point the split between Gomphocerinae and Oedipodinae, estimated to have occurred ∼100 Mya ago. This estimate is based on dated ancient cockroach fossils (Gaunt and Miles 2002, Fries et al. 2007). As a second calibration point, we used the divergence between *Sphingonotus azurescens* (mainland species) and S. *guanchus* (endemic to La Gomera Island, Canary Islands), whose estimated age is around 3.5 Mya (Husemann et al. 2014). *Sphingonotus guanchus* was only included in the BEAST analysis for calibration purposes. Substitution parameters were based on analyses previously conducted in jModeltest. We performed two independent runs of 100 × 10^5^ generations, sampled every 100,000 generations. We then used Tracer 1.4.1. to examine convergence, determine the effective sample sizes (EES) for each parameter, and compute the mean and 95% highest posterior density (HPD) interval for divergence times. We confirmed EES >200 was achieved for all parameters after the analyses. Tree and log files (12 000 trees after a 5% burn-in) of the two runs were combined with LogCombiner 1.4.7 (Drummond et al. 2012), and the maximum clade credibility (MCC) tree was compiled with TreeAnnotator 1.4.7. The concatenated matrix and the ML and Bayesian phylogenies (in Nexus format) are available in Dryad and TreeBASE Web, respectively. The obtained phylogenies were robust and largely consistent with previous studies (Nattier et al. 2011).

### Morphological data

Adult size was characterized from the length of the left hind femur. We used femur length as an indicator of body size since the total length of females varies substantially with the oviposition cycle (Hochkirch and Gröning 2008). Femur length was strongly correlated with structural body length excluding the abdomen (i.e. head + thorax) in both sexes (males: *R*^2^ = 0.70, *p* <0.001; females: *R*^2^ = 0.65, *p* <0.001). Femur length scales isometrically with body length (*β*_males_ = 0.964, *β*_females_ =1.048) and thus, constitutes a good proxy for adult body size (see also Ortego et al. 2012, Laiolo et al. 2013, Anichini et al. 2016, Bidau et al. 2016). Hind legs were carefully removed from the body of adults in the laboratory under a ZEISS stereomicroscope (SteREO Discovery V.8; Carl Zeiss Microscopy GmbH, Germany) and photographed using ZEISS image analysis software (ZEN2). Measurements of femur length were made on a total of 720 individuals, 365 males and 355 females (4-5 individuals of each sex per species). Because SSD is practically always female sex-biased in most orthopteroids, we quantified sexual size dimorphism as the ratio of female to male femur length (the simplest SSD estimator; see Lovich and Gibbons 1992). In addition, we quantified the relative length of the stridulatory file (i.e. length of the stridulatory file/femur length*100) for Gomphocerinae species. The stridulatory file consists of a row of pegs located on the inner side of the femur of each hind leg (e.g. Jago 1971). Gomphocerine grasshoppers produce acoustic signals (songs) by rubbing this structure against the forewings. Calling songs are used to search for conspecific mates and, thus, the evolution of the stridulatory apparatus is expected to be subject to sexual selection (von Helversen and von Helversen 1994, Mayer et al. 2010, Nattier et al. 2011). Males with larger sound-generating organs are able to produce low frequency sound, which is associated with larger male body size (see Anichini et al. 2016 and references therein). For example, in a comparative study of 58 bushcricket species, Montealegre-Z (2009) showed that the length of the stridulatory file correlated positively with male body size and pulse duration. Hence, we used the relative length of the stridulatory file as a proxy of the strength of pre-copulatory sexual selection in this subfamily.

### Life-history and ecological data

#### i) Fecundity

Given that large females can generally allocate more resources and energy to reproduction resulting in more offspring and/or higher-quality offspring, fecundity selection usually favors larger body size in females (reviewed in Pincheira-Donoso and Hunt 2016). We tested for fecundity selection by using mean ovariole number (which reflects the number of eggs produced in a discrete time interval) as an index of a species’ potential fecundity. Ovariole number is a strong determinant of fecundity, and therefore fitness because it sets the upper limit for reproductive potential (i.e. females with more ovarioles can produce more eggs in a discrete time interval) (Stauffer and Whitman 1997, Taylor and Whitman 2010). This parameter was extracted from different sources (Ingrisch and Köhler 1998, Reinhardt et al. 2005, Schultner et al. 2012) for a subset of species (*n* = 20; see Supporting Information).

#### ii) Substrate use

The substrate or structure on which a species rests can have major implications for the evolution of body size and SSD. For example, Moya-Laraño et al. (2002) proposed the so-called “gravity hypothesis” to explain patterns of SSD in spiders whereby species building webs high in the vegetation are predicted to show greater SSD than those that build lower down. Whilst, Brandt and Andrade (2007) proposed that the prevalence of female-biased SSD in species that occupy elevated substrates may be explained by selective advantages for small males in vertical habitats and for large males of low-dwelling species to run faster on the ground. In this sense, a previous study has shown that life form correlated with individual size in grasshoppers from Inner Mongolia (Yan and Chen 1997); larger species were typically terricoles, whereas the smaller ones were typically planticoles. Here, we tested if ground-dwelling grasshoppers exhibit a lower level of SSD than those species that perch on plants after correcting for phylogeny. To that end, each species was assigned to one of the two categories (ground *vs*. plant-perching) based on the literature (e.g. Default and Morichon 2015), personal observations and a survey of photographs available in open-access online repositories (http://www.biodiversidadvirtual.org; http://www.pyrgus.de; http://www.orthoptera.ch).

#### iii) Length of the breeding season

Season length has been postulated as another important factor in driving body size evolution. Individuals can become larger by lengthening the growth period but at the expense of a high cost: they may die before reproducing. In contrast, for example in ephemeral habitats, an individual can rush through its development in order to reach adulthood faster and reproduce (Roff 1980, Blanckenhorn and Demont 2004). In this sense, the length of the breeding season in conjunction with ambient temperature has been postulated as the main cause for the existence of altitudinal phenotypic clines in many ectothermic species with short generation times (Masaki 1967, Chown and Klok 2003). In Orthopterans, several studies have reported a reduction in body size with altitude (e.g. Berner and Blanckenhorn 2006, Bidau and Martí 2007a, but see also Sanabria-Urbán et al. 2015). Accordingly, we would expect small adult size in species with a short life-span and/or species inhabiting higher altitudes (supposedly more seasonal habitats) because a shorter growing season should select for earlier maturation and smaller body size. To test such a hypothesis, we compiled information on the length (in months) of the life-cycle of each species from the available literature (e.g. Lluciá-Pomares 2002) and our own field observations. The length of the breeding season oscillated between 2 and 12 months, that is, from species only present as adults during the summer period to those present all year round. Species that have adults present year round likely have more than one generation each year (i.e. bivoltine species) and thus a period of sexual diapause. Because it might compromise the reliability of our results, we repeated our analyses after excluding such species (five excluded species). In addition, a subset of species was classified into three categories (low-altitude, medium-altitude, high-altitude) according to the altitudinal range in which these species can be found (e.g. Pardo and Gómez 1995; Lluciá-Pomares 2002; authors, *pers. obs*.). Those species (*n* = 16) with a broad altitudinal range (e.g. from 0 to 2000 m; see Supporting Information) were discarded from this analysis.

#### iv) Habitat specialization

The level of ecological specialization of an organism, that is, its variance in performance across a range of environmental conditions or resources, has implications in terms of population density and local competition (e.g. Devictor et al. 2010, Parent et al. 2014), two factors often associated with the extent of sexual dimorphism. Selection for larger male size is expected to be greater in species with a narrow ecological niche (i.e. specialist species) and/or limited dispersal ability due to strong intrasexual competition for resources. Accordingly, we predict higher levels of SSD in generalist species. In order to obtain a measure of ecological specialization, we calculated the so-called ‘Paired Difference Index’ (PDI) (Poisot et al. 2011, 2012) from a species-habitat matrix in which we rated the level of association of each species (from complete generalist, 0, to complete specialist, 3) with the nine most common habitats in which these species can be found (see Supporting Information). The PDI is a robust specialization index which takes into account not only the number of resources used by a species, but also the strength of the association between the species and its resources (Poisot et al. 2012). Scores of species-habitat association were obtained directly from the literature (research articles, monographs and field guides) and our own personal observations (Table S1, Supporting Information). PDI values were computed using the *bipartite* package in R (Dormann et al. 2016).

### Phylogenetic comparative analyses

For each studied variable (male size, female size, SSD, length of the breeding season, length of the stridulatory file, ovariole number, PDI) we assessed the amount of phylogenetic signal, a measure of how similar closely related species are to one another for a given trait. To assess phylogenetic signal we used Pagel’s lambda (λ; Pagel 1999) and Blomberg’s *K* (Blomberg et al. 2003) computed using the “phylosig” function in the R package *phytools* (Revell 2011). To visualize substrate (binary variable; ground: 0; plant-perching: 1) variation among species on the phylogentic tree we used maximum likelihood reconstruction in MESQUITE v. 3.04 (Maddison and Maddison 2015). We also reconstructed ancestral states for our focal trait, SSD, in Mesquite.

We tested for departure from a null model of constant rate of diversification using the γ statistic as implemented in *ape* (Paradis et al. 2014). A significantly negative value of γ indicates a deccelerating rate of cladogenesis through time. The γ statistic is biased by incomplete taxon sampling, because the number of divergence events tends to be increasingly underestimated toward the present (favoring negative values for the γ). Therefore, we corrected for undersampling using the Markov chain constant-rates (MCCR) test (Pybus & Harvey 2000) as implemented in the *laser* package (Rabosky 2006). Recent MEDUSA analyses performed by Song et al. (2015) indicate that the lineage leading up to Acrididae has undergone a significant increase in diversification rate with little or no extinction. Thus, values of γ are unlikely to be biased by extinction rates.

We investigated the mode of male and female body size evolution by comparing fits of these traits to four different models of evolution using the Akaike information criterion (Akaike weights, AICw, and size-corrected Akaike values, AICc): i) pure Brownian Motion (null) model (BM), ii) Ornstein-Uhlenbeck (OU), iii) ‘early-burst’ (EB) and iv) time-variant rate (TVR) model. Under BM, traits evolve along a random walk whereby each change is independent of the previous change (morphological drift; Felsenstein 1985). The OU model describes a random walk with a single stationary peak, such that trait values have a tendency to return to an optimal value (θ) (Hansen 1997, Butler and King 2004). Under an EB model, the net rate of evolution slows exponentially through time as the radiation proceeds (Blomberg et al. 2003), whereas the TVR model is similar to an EB model but also allows an exponential increase of evolutionary rates through time (Pagel 1999). The TVR model can be used to evaluate the non-constant rate of evolution through time using the path-length scaling parameter Pagel’s delta, δ. This parameter detects differential rates of evolution over time (i.e. δ = 1 means gradual evolution).

If δ <1, shorter paths contribute disproportionately to trait evolution (decelerating evolution) whereas if δ >1 is the signature of accelerating evolution as time progresses (see Hernández et al. 2013). Specifically, we expected female size to show a trend towards larger sizes (i.e. directional selection for increased female size) whereas males would likely be maintained at optimal values (i.e. directional selection for the maintenance of small male size according to an OU model). From these models we calculated the evolutionary rate (σ^2^) for each sex in order to determine whether body size evolution of males was faster or slower than body size evolution of females. Evolutionary models were run using the R package *geiger* (Harmon et al. 2008).

Additionally, we applied two complementary methods: the morphological diversity index (MDI, Harmon et al. 2003) and the node-height test (Freckleton & Harvey 2006) to analyze patterns of evolution. Both methods test for departure from Brownian motion but differ in the approach used to test for this departure. First, we calculated disparity through time (DTT) plots using the ‘dtt’ function in the *geiger* package. DTT analyses compare phenotypic diversity simulated under a Brownian Motion model with observed phenotypic diversity among and within subclades relative to total disparity at all time steps in a time-calibrated phylogeny. Low (i.e. negative) values of relative disparity indicate that most morphological disparity originated early in the history of the group (early divergence), whereas high (positive) values indicate that most morphological disparity originated more recently compared to a random walk pattern (recent phenotypic divergence). Values near 0 indicate that evolution has followed BM. The MDI was calculated as the sum of the areas between the curve describing the morphological disparity of the trait and the curve describing the disparity under the null hypothesis of BM (1 000 simulations). Finally, we used the node-height test (Freckleton & Harvey 2006) to test whether grasshopper body size evolution has slowed over time. We computed the absolute value of standardized independent contrasts for body size on our MCC tree and correlated them with the height of the node at which they are generated. A significant negative relationship between absolute contrast value and node age implies that rates of body size evolution slow down over time according to a niche-filling model (‘early-burst’ of trait evolution).

### Ecological correlates of body size and SSD

To explore the association between SSD and our continuous ecological (habitat specialization, breeding season length) and reproductive (fecundity, length of the stridulatory file) variables, we used phylogenetic generalized least squares (PGLS_λ_). Maximum likelihood estimates of the branch length parameters delta (a measure of disparity of rates of evolution through time, see above), lambda (a measure of phylogenetic signal, see above) and kappa (which contrasts punctuational *vs*. gradual trait evolution, see Hernández et al. 2013) were obtained to optimize the error structure of the residuals in each comparison as recommended by Revell (2010). PGLS regression analyses were performed using the R package *caper* (Orme 2013) and graphically visualized by means of phylogenetically independent contrasts (PIC) computed using the PDAP:PDTREE module in Mesquite (Midford et al. 2005). To test the influence of categorical variables (substrate, altitude class) on our focal traits independently from the phylogeny, we employed phylANOVA (10 000 simulations) as implemented in the R package *phytools* (Revell 2011).

We tested for greater evolutionary divergence in male size compared with female size (Rensch’s rule test) by regressing log-transformed male body size against log female body size using phylogenetic major axis regressions (PRMA; Revell 2011), a method that accounts for the shared evolutionary history of species. This analysis was performed at two taxonomic levels, across our entire phylogeny and within the two largest subfamilies (Gomphocerinae, Oedipodinae) because Rensch’s rule was originally proposed for ‘closely-related species’ (Rensch 1950). We tested if the slope (*β*) of the regression of body size of males on females was larger than 1 (as predicted by Rensch’s rule), smaller than 1 (converse of Rensch’ rule) or not different from 1 (i.e. *β* = 1; isometric pattern) (see Ceballos et al. 2013 for more details about the possible scenarios for the relationship between SSD and body size of males and females).

Statistical significance of the allometric pattern was determined based on the 95% confidence intervals (CI) of *β* using the *smatr* R package (Warton et al. 2012).

## Results

### Phylogenetic signal

Both male and female body size exhibited a strong phylogenetic signal (male body size: λ = 0.955, *p* <0.01; *K* = 0.267, *p* = 0.016; female body size: λ = 0.956, *p* <0.001; *K* = 0.213, *p* = 0.03), which indicates that the body size of related species is more similar than expected under Brownian Motion. Accordingly, we also found a strong phylogenetic signal for SSD (λ = 0.904, *p* = 0.03; *K* = 0.225, *p* <0.01; Fig. 1). The relative length of the stridulatory file showed a moderate phylogenetic signal (λ = 0.589, *p* = 0.09; *K* = 0.114, *p* = 0.02), whereas the level of ecological specialization (PDI) (λ ∼ 0, *p* = 1; *K* = 0.06, *p* = 0.36) and the length of the breeding season (λ = 0.627, *p* = 0.01; *K* = 0.107, *p* = 0.08; Fig. 1) did not show phylogenetic inertia. Ovariole number showed a strong phylogenetic signal (λ ∼ 1, *p* <0.001; *K* = 1.987, *p* <0.001). Substrate type (ground *vs*. plant-perching) seems to be a conservative trait in short-horned grasshoppers; ground-species are predominant in the Oedipodinae subfamily whereas plant-perching species are more abundant within the Gomphocerinae subfamily (see Fig. 1).

**Fig. 1.**
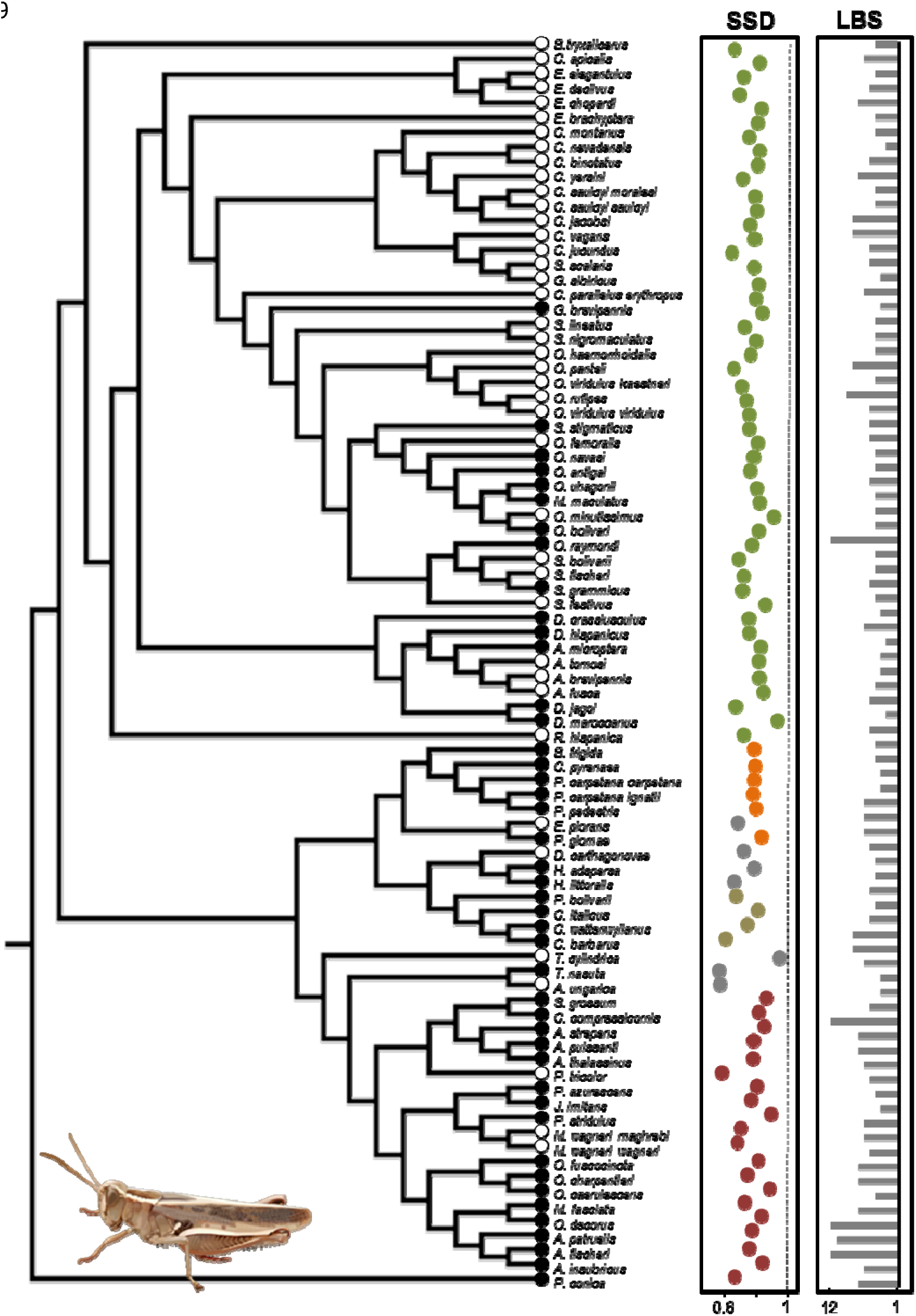
Variation in degree of sexual size dimorphism (SSD) and length of the breeding season (LBS, in months) among the studied Acrididae species (in the SSD scatterplot species belonging to the same subfamily are indicated with the same color). Color of dots next to tips in the tree denotes the main substrate used by each species (black dots: ground; white dots: plant).

### Tempo and mode of body size evolution

The rate of diversification accelerated with time (γ = 1.68) indicating that rapid diversification occurred late in the evolutionary history of the group. When comparing alternative models of evolution across the entire dataset, the OU model (Brownian Motion with selective constraint) exhibited the best fit for the evolution of male body size whereas the OU and the TVR models were equally supported for female body size (ΔAICc <2) (Table 1). When restricting our analyses to the Gomphocerinae subfamily, we found that the best-supported model for the evolution of both male and female body size was TVR (Table 1), suggesting that trait evolution is non-constant through time. In Oedipodinae, the BM model provided a better fit than the other models for the evolution of male body size whereas BM and the two non-constant models (EB and TVR) were similarly supported (ΔAICc <2) in females (Table 1). The comparison of evolutionary rates between sexes indicated that body size evolved at a similar pace (Table 1).

**Table 1.**
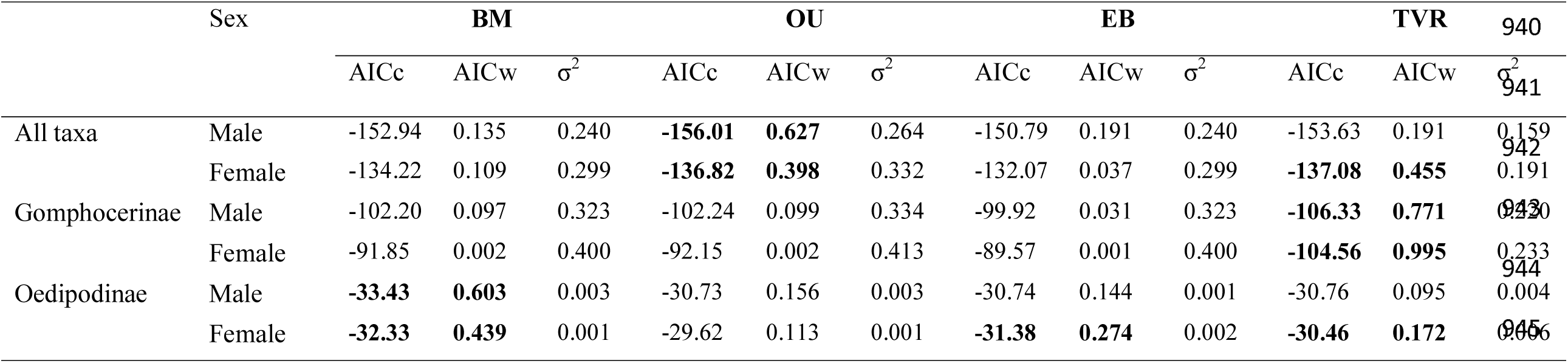
Relative support for alternative evolutionary models of male and female body size in short-horned grasshoppers. BM: Brownian Motion; OU: Ornstein-Uhlenbeck; EB: early-burst; TVR: time-variant rate model. σ^2^ denotes the estimated reproductive rate for each sex. AICc, corrected Akaike’s information criterion (AIC) value; AICw, AICc weight. The best-fit model/s is/are highlighted in bold type.

|  |  |  |  |  |  |  |  |  |  |  |  |  | 939 |
| --- | --- | --- | --- | --- | --- | --- | --- | --- | --- | --- | --- | --- | --- |
|  | Sex | BM |  |  | OU |  |  | EB |  |  | TVR |  | 940 |
| | | AICc | AICw | $\sigma^2$ | AICc | AICw | $\sigma^2$ | AICc | AICw | $\sigma^2$ | AICc | AICw | $\sigma^2$ |
| All taxa | Male | -152.94 | 0.135 | 0.240 | **-156.01** | **0.627** | 0.264 | -150.79 | 0.191 | 0.240 | -153.63 | 0.191 | 0.159 |
|  | Female | -134.22 | 0.109 | 0.299 | **-136.82** | **0.398** | 0.332 | -132.07 | 0.037 | 0.299 | **-137.08** | **0.455** | 0.191 |
| Gomphocerinae | Male | -102.20 | 0.097 | 0.323 | -102.24 | 0.099 | 0.334 | -99.92 | 0.031 | 0.323 | **-106.33** | **0.771** | 0.130 |
|  | Female | -91.85 | 0.002 | 0.400 | -92.15 | 0.002 | 0.413 | -89.57 | 0.001 | 0.400 | **-104.56** | **0.995** | 0.233 |
| Oedipodinae | Male | **-33.43** | **0.603** | 0.003 | -30.73 | 0.156 | 0.003 | -30.74 | 0.144 | 0.001 | -30.76 | 0.095 | 0.004 |
|  | Female | **-32.33** | **0.439** | 0.001 | -29.62 | 0.113 | 0.001 | **-31.38** | **0.274** | 0.002 | **-30.46** | **0.172** | 0.006 |

Maximum likelihood estimates of δ computed for all taxa and for each of the two subfamilies were high (δ values: all clades; male body size: 2.45, female body size: 3.00; *Gomphocerinae;* male body size: 2.53, female body size: 3.00; *Oedipodinae;* male body size: 3.00, female body size: 3.00) suggesting that longer paths (i.e. later evolution of the trait in the phylogeny) contribute disproportionately to trait evolution (‘late-burst’).

DTT plots showed that phenotypic disparity within lineages is greater than expected under a BM model late in the diversification of Acrididae (Fig. 2). We obtained positive MDI statistics, indicating that the proportion of total morphological disparity within clades was less than expected by a Brownian Motion model for all taxa and each of the two subfamilies (average MDI values: all clades; male body size: 0.138, female body size: 0.101; *Gomphocerinae*; male body size: 0.342, female body size: 0.212; *Oedipodinae*; male body size: 0.088, female body size: 0.357).

**Fig. 2.**
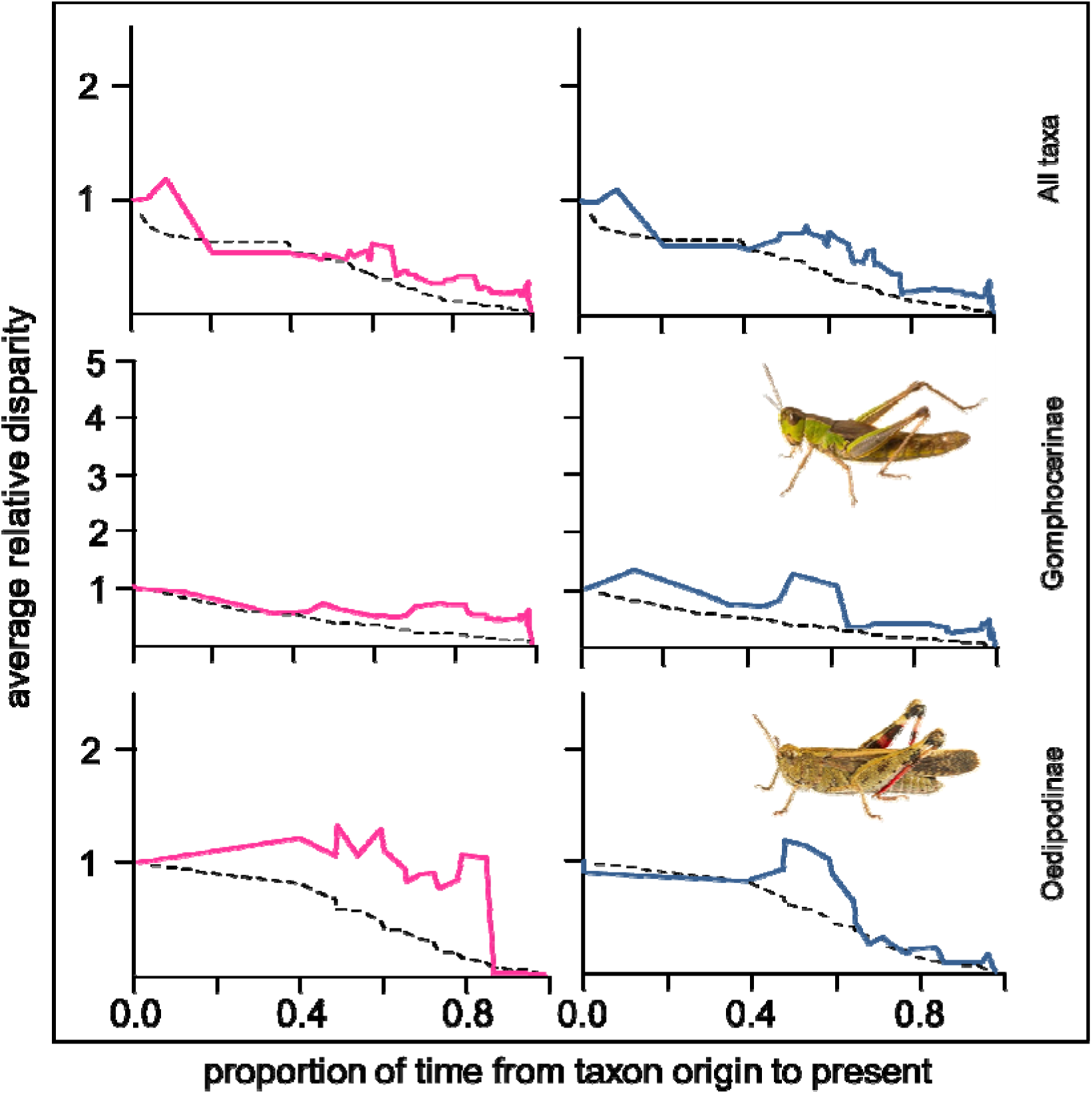
Relative disparity plots for short-horned grasshoppers compared with expected disparity based on phylogenetic simulations. The continuous line shows the actual pattern of phenotypic disparity (graphs are color-coded according to sex; pink color: female body size, blue color: male body size) and the dotted line represents the median result of Brownian model simulations (1 000 simulations). Time is relative to phylogenetic depth from the base of the phylogeny on the left to the terminal tips on the right.

Concordant with late shifts in the acceleration of the net diversification rate, the node-height test resulted in a positive but non-significant relationship between the absolute values of standardized length contrasts and node age in both sexes (*males: t* = 1.01, d.f. = 83, *p* = 0.32; *females: t* = 0.72, d.f. = 83, *p* = 0.47) across all taxa. For the Gomphocerinae subset, we found a positive and significant relationship between the absolute value of independent contrasts and the height of the node from which they were generated (*males: t* = 2.46, d.f. = 45, *p* = 0.02; *females: t* = 3.11, d.f. = 45, *p* = 0.003), indicating that body size evolution has increased through time. When restricting our analyses to the Oedipodinae subfamily, the node height test yielded a negative and non-significant relationship for the evolution of male and female body size (*male body size: t* = -0.60, d.f. = 18, *p* = 0.55; *female body size: t* = -1.02, d.f. = 18, *p* = 0.32).

### Ecological correlates of body size and SSD

We found a negative significant association between SSD and female body size (PGLS; estimate: -0.444 ± 0.102, *t* = -4.34, *p* <0.001) but not such association between SSD and male body size (PGLS; *t* = 1.05, *p* = 0.30) across taxa. A similar result was found for the Gomphocerinae subset (PGLS, *female size:* estimate: -0.458 ± 0.101, *t* = 4.492, *p* <0.001; *male size: t* = 0.52, *p* = 0.60). In contrast, in Oedipodinae, we obtained the opposite pattern: a significant association between SSD and male body size (PGLS; estimate: 0.666 ± 0.138, *t* = 4.82, *p* <0.001) but not between SSD and female body size (PGLS; *t* = 0.054, *p* = 0.95). Overall, the emerging picture was that both sexes tend to be progressively more similar as size increases. When size decreases within one lineage through evolutionary time, then male size decrease disproportionately with respect to female body size.

When testing for fecundity selection, we failed to find a significant relationship between ovariole number (a good proxy for the reproductive potential of a given species) and female body size (PGLS; *n* = 20, *t* = 0.067, *p* = 0.947). When only considering gomphocerine grasshoppers, no significant association was found between the size-corrected length of the stridulatory file and SSD (PGLS; *n* = 48, *t* = 0.12, *p* = 0.90) (Fig. 3). Short-lived species did not show either a greater degree of development of the stridulatory organ as expected if the strength of sexual selection is higher in species with a shorter breeding period (PGLS; *n* = 48, *t* = -0.59, *p* = 0.55). The degree of SSD was phylogenetically correlated with the length of the breeding season; species with a long phenology are more dimorphic in size that those with a short reproductive window (PGLS; estimate: -0.010 ± 0.004, *n* = 85, *t* = -2.16, *p* = 0.035; Fig. 1).

**Fig. 3.**
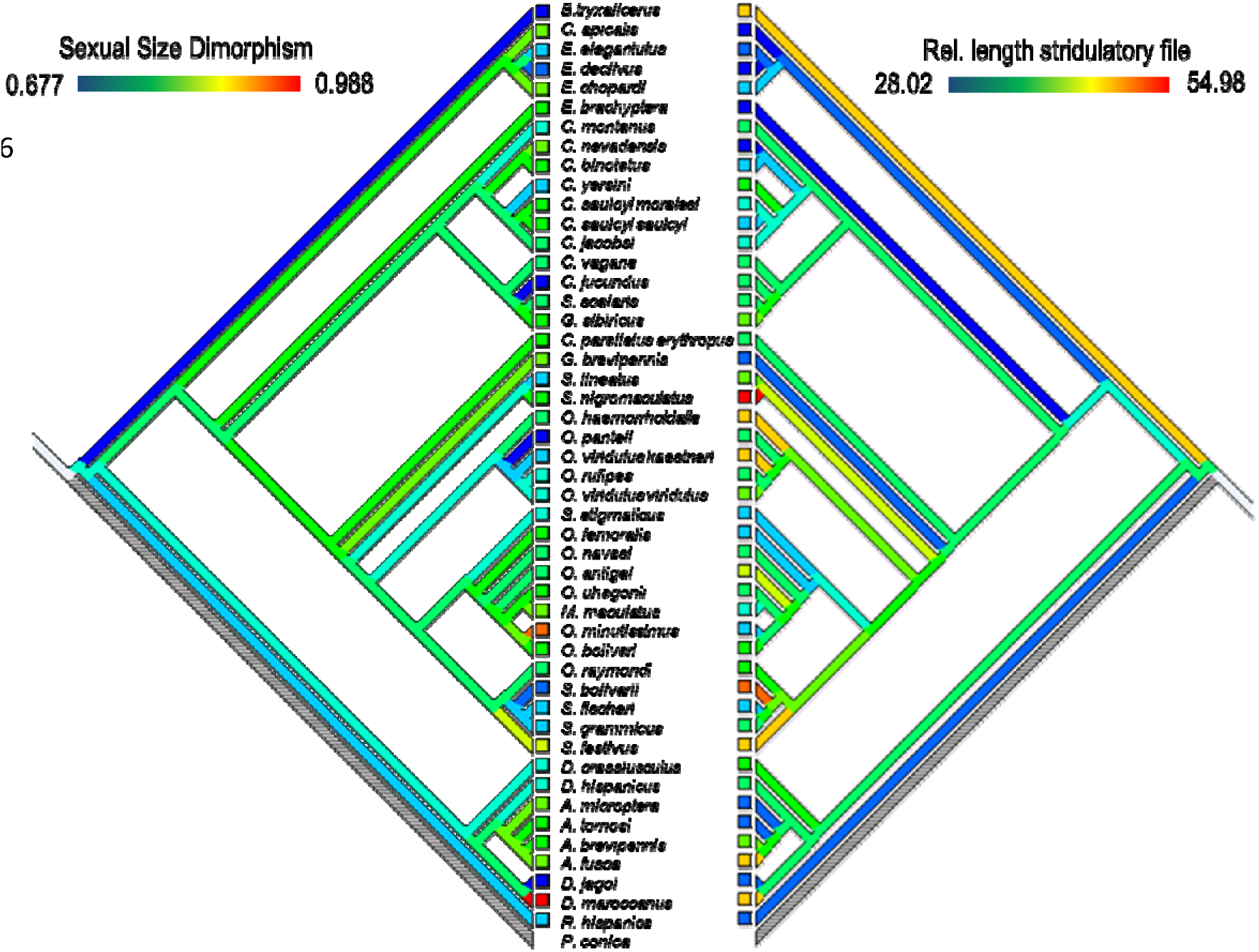
Reconstructed evolution of sexual size dimorphism (sexual size dimorphism ratio, with lower values indicating more dimorphic species and higher values indicating less dimorphic species, so 1 denotes monomorphism) and relative (size-corrected) length of the stridulatory file (length of the stridulatory file/femur length*100) in the grasshopper subfamily Gomphocerinae. Colors denote size classes (see legends).

Such a relationship became more significant when excluding year-round species (length of the breeding season = 11-12 months) from the dataset (PGLS; estimate: -0.011 ± 0.004, *n* = 80, *t* = - 2.34, *p* = 0.021; Fig. 4). Although we found that habitat specialist species (high PDI values) have a shorter breeding cycle than more generalist species (PGLS, estimate: -6.596 ± 1.545, *n* = 85, *t* = -4.27, *p* <0.001), there was no significant relationship between the level of SSD and the degree of ecological specialization (PDI index) (PGLS; *t* = -1.16, *p* = 0.25).

**Fig.4.**
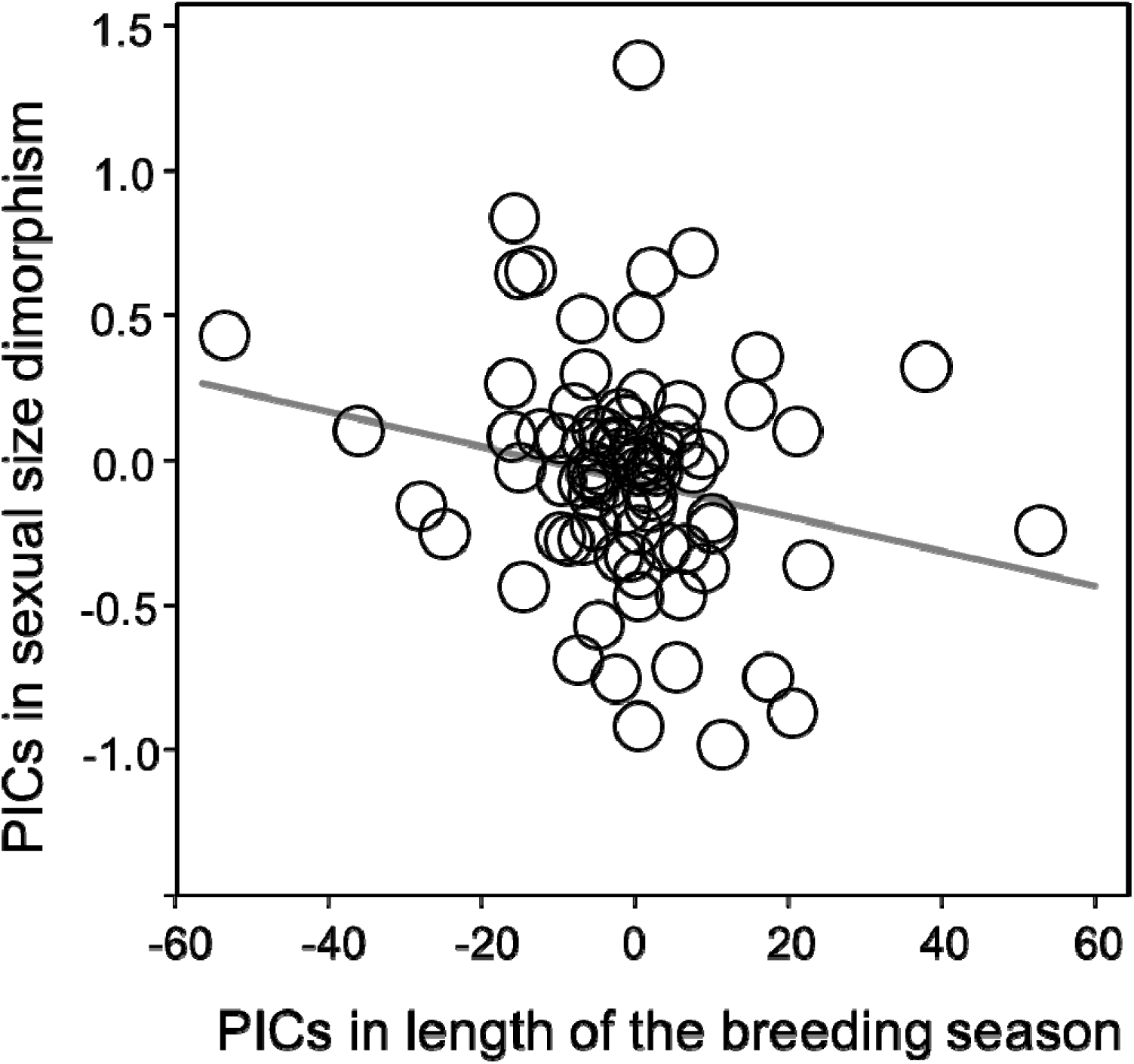
Relationship between sexual size dimorphism and the length of the breeding season represented in the form of standardised phylogenetic independent contrasts (PICs).

Regarding categorical variables, we did not find significant differences between ground and plant-perching species in terms of SSD (PhylANOVA; F_1,83_ = 2.72, *p* = 0.51), male body size (F_1,83_ = 0.69, *p* = 0.75) or female body size (F_1,83_ = 0.02, *p* = 0.96) after controlling for the shared evolutionary history of species. Lastly, there was a trend towards low altitude species being larger than those inhabiting medium- and high-altitudes, but differences were not statistically significant after correcting for phylogeny (PhylANOVA; *female body size*: *F*_2,66_ = 8.51, *p* = 0.13; *male body size:* F_2,66_ = 6.38, *p* = 0.21) (Fig. 5). The length of the breeding season, but not the level of SSD, differed significantly among altitude categories (breeding season length: *F*_2,66_ = 4.22, *p* = 0.018; SSD: *F*_2,66_ = 3.23, *p* = 0.45), being shorter at higher altitudes (mean ± S.D. breeding season length; low-altitude: 5.83 ± 1.01, medium-altitude: 5.36 ± 2.46, high-altitude: 4.07 ± 2.70 months).

**Fig. 5.**
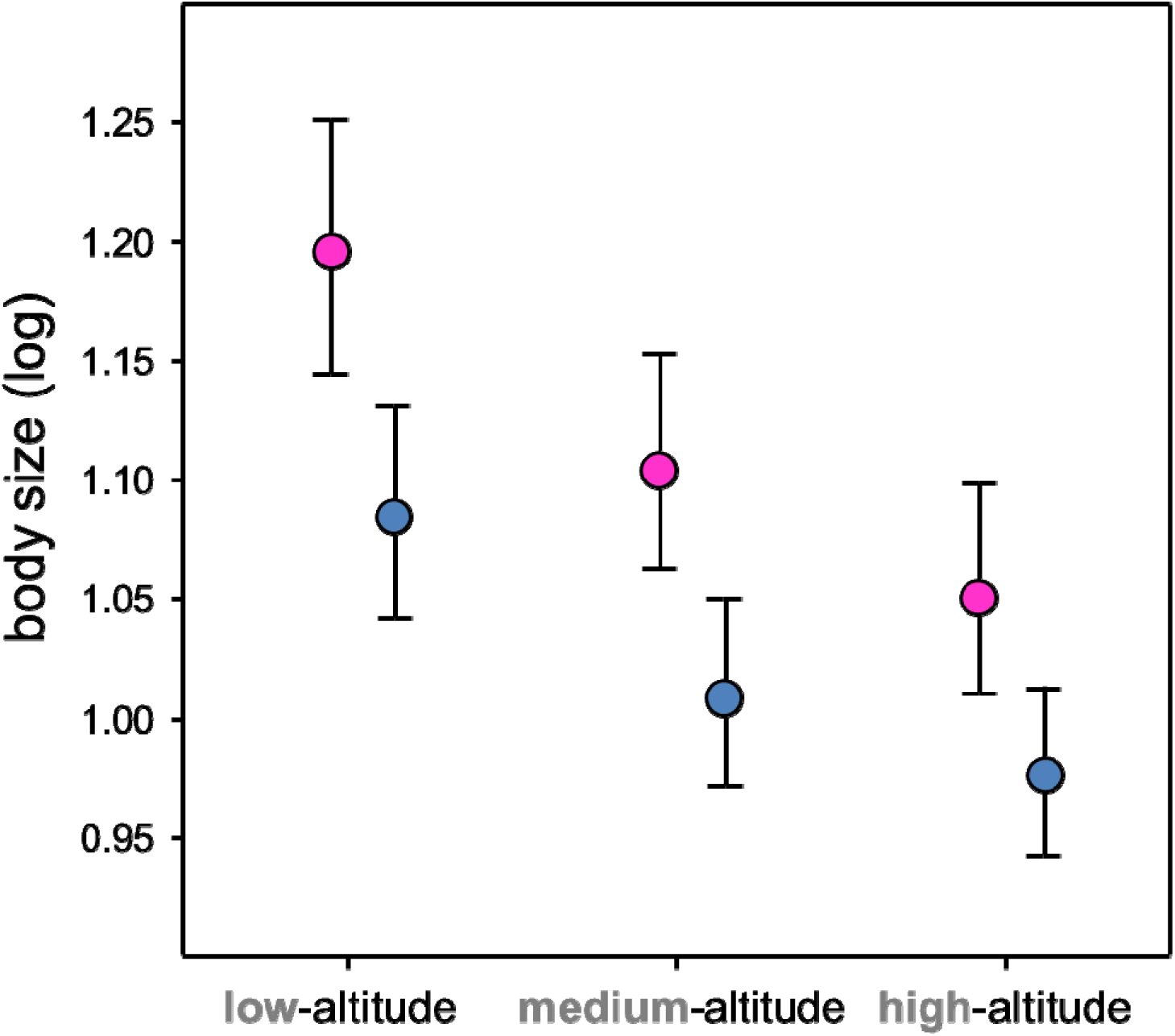
Differences in average male (blue dots) and female (pink dots) body size between grasshopper species inhabiting low (<800 m a.s.l., *n* = 18), medium (800-1500 m, *n* = 25) and high-altitudes (>1500 m, *n* = 26) (*n* = 18, 25, and 26, respectively).

### Allometry of SSD: Do Acrididae grasshoppers conform to Rensch’s rule?

The degree of SSD was similar across all taxa (all taxa: 0.801 ± 0.076) and when considering the two most speciose subfamilies separately (Gomphocerinae: 0.808 ± 0.062; Oedipodinae: 0.817 ± 0.076). Our results supported the existence of a pattern consistent with the converse to Rensch’s rule as the relationship of male body size with female body size across all taxa had a slope less than one (PRMA; *β* = 0.895, CI: 0.804-0.940). However, we found an isometric pattern (i.e. the slope did not differ significantly from 1) when data were analyzed separately for Gomphocerinae (PRMA; *β* = 0.898, CI: 0.869-1.106) and Oedipodinae (PRMA; *β* = 0.866, CI: 0.846-1.422).

## Discussion

### Evolution of body size in a recent radiation

The very short branches deep in our phylogeny suggest recent divergence and thus rapid speciation in some lineages. This is in agreement with the findings of Song et al. (2015), who suggested that Acrididae may have undergone an explosive adaptive radiation during the Cenozoic, when global climate became temperate and grasses evolved and became dominant.

The evolution of a new niche space (grasslands) may have powered the radiation of graminivorous species, especially strong fliers like band-winged grasshoppers (Oedipodinae) (Song et al. 2015). Later, climatic oscillations during the Pleistocene led to thermophilic species (as most Gomphocerinae are) being restricted to southern refuges during glacial stadials. This probably resulted in divergent evolution of allopatric populations by geographic isolation (Taberlet et al. 1998, Hewitt 1999, Mayer et al. 2010). This plausible scenario matches with the notion raised by Schluter (2000) who noted that “a continuous spread to new environments is the dominant trend of adaptive radiation”. Although the term adaptive radiation is frequently used to describe a slowdown in diversification and morphological evolution after an initial phase of rapid adaptation to vacant ecological niches (‘niche filling’), recent studies stress that the definition of adaptive radiation should not be conditioned by the existence of early-bursts, which indeed seem to be uncommon across the tree of life (Harmon et al. 2010, Pincheira-Donoso et al. 2015). Rather, an adaptive radiation should be defined as the process in which a single lineage diversifies into a variety of species, occuring at a fast net rate, irrespective of the timing (Harmon et al. 2010). In this study, model comparison, maximum-likelihood values for the δ parameter (which tests for acceleration vs. deceleration) and node-height tests provide no significant support for an ‘early burst’ followed by a slowdown in morphological evolution in this taxonomic group. Instead, we found the opposite pattern; the high values of δ indicate recent, high rates of phenotypic divergence whereas the results of the node-height tests indicate that it increased as the number of taxa increased. This pattern suggests that most divergence seems to be concentrated later in the evolutionary history of this group (i.e. recent and rapid diversification; see also Boucher et al. 2012, Edwards et al. 2015).

Body size evolution in Iberian Acrididae is inconsistent with a Brownian Motion process, indicating that selection and not drift underlies body size evolution in this group. Our results indicate that body size variation is best explained by a time-dependent model (TVR model; Pagel 1997, 1999). However, across all subfamilies the OU model provided a better fit to the data suggesting a process of stabilizing selection in which variation of body size revolves around stationary optimal values. That is, deviant body sizes are “polished” towards an optimum value, which was estimated to be around 9.79 and 13.49 mm for male and female femur length, respectively. The pattern described for the entire family reflects the existence of scale-dependent processes that act differentially across stages of the diversification process (see Ceballos et al. 2013). Overall, this evidences that even in taxonomic groups showing limited morphological and ecological disparity, natural selection seems to play a more important role than genetic drift in driving the radiation process.

### Ecological correlates of body size and SSD

Although the predominance of female-biased SSD in Acrididae suggests that fecundity selection may be the most important selective force acting on this family, we failed to find support for this hypothesis as ovariole number and female body size were not correlated. However, this specific analysis was performed using a reduced dataset (*n* = 20) and thus, our results should be interpreted cautiously. Alternatively, it is also likely that fecundity selection acts primarily at intraspecific level, preventing us to detect its effect through our analyses. Regarding sexual selection, we failed to find a significant correlation between the relative length of the stridulatory file (our proxy to measure the strength of sexual selection) and the level of SSD. This may indicate that sexual selection is not driving SSD in Gomphocerinae or, alternatively, that this trait does not accurately reflect the strength of this selective force at interspecific level due to its strong genetic component (Saldamando et al. 2005). The size-corrected length of the stridulatory apparatus showed a non-phylogenetic signal, supporting the view that in gomphocerine grasshoppers the value of acoustic characters as an indicator of phylogeny is very limited despite the fact that they have long been used to resolve taxonomic uncertainties at the species level (Ragge 1987, Ragge and Reynolds 1988).

Substrate use (ground or plant-perching) may be expected to affect body size as different functional demands between the two types are expected to generate different selective peaks. However, we did not find evidence for the gravity hypothesis; ground-dwelling species did not show a significantly larger size than plant-perching species as expected if climbing ability selects for reduced body size (Moya-Laraño et al. 2012). Thus, we failed to detect strong selection for increased female size (fecundity selection) (see above) or for the maintenance of small male size (agility hypothesis) in these insects. Regarding the other ecological variable, habitat specialization (PDI index), we did not find a higher level of SSD in generalist species in which selection for large male body size should be smaller in comparison with species with specific habitat requirements and whose populations may be affected by higher intrasexual competition. A plausible explanation for this result is that male-male competition for food resources may not be intense enough to boost the evolution of male body size as most species depend on food resources that are rarely limited (e.g. gramineous). This finding thus reinforces the view that both inter- and intrasexual competition for food is unlikely in small herbivorous organisms and that these play a subsidiary role in the evolution of SSD (Fairbarn et al. 2007).

Larger-bodied and slower developing arthropods like Orthoptera are expected to be more affected by seasonal limitations than faster developing insects. Shorter breeding season lengths should promote life history adaptations leading to smaller body size to facilitate the completion of the life cycle within the reduced time available for development. When the developmental time window is short, individuals reach maturity at smaller sizes and develop faster. When the length of the breeding season is longer, more time is available to reach the reproductive stage at a larger size (Berner and Blanckenhorn 2006). Contrary to our initial expectation, we found a lower level of SSD in species with a short phenology. SSD was more pronounced in species with longer breeding seasons, an effect that seemed to be caused by a larger difference in size between sexes. It suggests that selection pressure for large body size in males may be stronger in ephemeral or more seasonal environments (i.e. in species with a short reproductive window). Alternatively, this result may be due to the fact that in most grasshopper species the females are more variable in size than males (for example, female size variation is exceptionally high in *Calliptamus* species), and that unfavorable environmental conditions may compromise body size (Teder and Tammaru 2005). This indicates that time constraints do not seem to impose limits for the evolution of male body size (rather the contrary, it seems to be favored) and that female body size is more sensitive to environment than male size. On the other hand, our results are consistent with the converse Bergmann’s rule; grasshopper body size tended to decrease with elevation but the differences were not statistically significant after correcting for phylogeny. This pattern is normally explained by the existence of gradients of precipitation and sun exposure, which are likely indicators of other ecological factors that exert control on body growth, such as resource availability and conditions for effective thermoregulation (Laiolo et al. 2013). Althought most evidence comes from single-species studies (Blanckenhorn et al. 2006; Bidau & Martí 2007b, 2008) and thus, our study is one of the first to test for altitudinal clines in body size at the interspecific level (that is, across species) in Orthoptera.

### Rensch’s rule

We found evidence that SSD and body size in short-horned grasshoppers fitted a converse Rensch’s rule: females are proportionally bigger than males in large species. This result is in agreement with previous studies carried out on a smaller scale (Bidau et al. 2013, Laiolo et al. 2013) and reinforces the view that Rensch’s rule is infrequent in taxononomic groups exhibiting female-biased SSD. When performing our analyses within subfamilies, we neither found a pattern consistent with Rensch’s rule: sizes of males and females scaled isometrically. A plausible explanation for these results is that if females are more sensitive to environmental conditions than males, they could achieve a better body size development under more bening conditions leading to an increase in SSD (Shine 1990, Stilwell et al. 2010). Thereby, Rensch’s rule and its converse would mirror sex-specific environmental sensibility and thus, these patterns may be considered subproducts of body size variations in relation to ecological conditions. Thereby, our study supports the idea that the so-called Rensch’s rule probably does not deserve the attribute “rule” at least in arthropods, wherein support for this pattern remains rather mixed (reviewed in Blanckenhorn et al. 2007a).

## Conclusions

Different and complex evolutionary pressures can affect body size evolution in Orthoptera. Fecundity, sexual selection, and predictable, long breeding season environments are thought to select for larger size, whereas time constraints, predators and unpredictable and poor-resource habitats are thought to select for small body size. Here, we found no support for either the fecundity or the sexual selection hypothesis, the two primary adaptive forces traditionally invoked to explain SSD. Nor did we find an effect of substrate -ground *vs*. plant- on body size evolution, a factor (agility) that has been suggested to explain why males of certain insect groups remain small. Our results also reinforce the idea that Rench’s rule is probably not a rule at all but a limited pattern only found in a few taxonomic groups and more frequently, at the intraspecific level (e.g. De Lisle and Rowe 2013, Liao et al. 2013, Bidau et al. 2016). Finally, and contrary to expected, we found a higher level of SSD in species with a long reproductive window, which is counter to the idea that SSD is favored in short-season habitats due to the fact that males have no time to fully develop (resulting in small adult sizes). These findings support laboratory studies at the intraspecific level showing that under poor conditions female Orthoptera are more strongly affected than males, reducing SSD (Teder and Tammaru 2005). We conclude that it is unlikely that protandry constitutes the main factor determining the existence of female-biased SSD in this insect radiation.

## SUPPORTING INFORMATION FOR

### Ecological drivers of body size evolution and sexual size dimorphism in short-horned grasshoppers (Orthoptera: Acrididae)

**Table S1.**Information on body size (male and female femur length, mm), length of the breeding season (months), substrate and altitude range for Iberian short-horned grasshoppers (Acrididae).

**Table S2.** Habitat requirements and ‘Paired Difference Index’ (PDI) values computed for the 85 grasshopper species included in this study.

| Taxa | Male size | Female size | Breeding season length |  | Substrate | Altitude range |
| --- | --- | --- | --- | --- | --- | --- |
| *Acrida ungarica mediterranea* | 22.40 | 37.33 | Ago-Oct | 3 | Plant | 0-900 |
| *Acrotylus insubricus* | 9.37 | 10.72 | Jan-Dec | 12 | Ground | 700-1500 |
| *Acrotylus fischeri* | 8.39 | 10.64 | Feb-Dec | 11 | Ground | 700-1800 |
| *Acrotylus patruelis* | 9.01 | 11.18 | Jan-Dec | 12 | Ground | 0-700 |
| *Aiolopus puissanti* | 10.70 | 13.10 | May-Nov | 7 | Ground | 0-700 |
| *Aiolopus strepens* | 11.07 | 12.50 | Jan-Dec | 12 | Ground | 0-1300 |
| *Aiolopus thalassinus* | 9.63 | 11.83 | May-Nov | 7 | Ground | 0-700 |
| *Arcyptera fusca* | 16.39 | 18.62 | Jul-Oct | 4 | Ground | 1300-2500 |
| *Arcyptera microptera* | 14.50 | 16.85 | Jun-Jul | 2 | Ground | 0-300 |
| *Arcyptera tornosi* | 14.92 | 17.58 | Jun-Ago | 3 | Plant | 1000-1900 |
| *Arcyptera brevipennis* | 13.50 | 15.84 | Jun-Ago | 3 | Plant | 800-1600 |
| *Bohemanella frigida* | 8.78 | 10.71 | Jul-Oct | 4 | Ground | 2000-3000 |
| *Brachycrotaphus tryxalicerus* | 8.60 | 12.36 | Jul-Oct | 4 | Plant | 0-800 |
| *Calephorus compressicornis* | 8.32 | 9.80 | Jun-Oct | 5 | Ground | 0-800 |
| *Calliptamus barbarus* | 8.84 | 13.88 | May-Nov | 8 | Ground | 0-2000 |
| *Calliptamus italicus* | 11.50 | 13.65 | Jul-Oct | 4 | Ground | 200-1600 |
| *Calliptamus wattenwylanus* | 11.20 | 14.40 | Jun-Oct | 5 | Ground | 600-1100 |
| *Chorthippus apicalis* | 11.28 | 13.20 | Apr-Sep | 6 | Plant | 0-1100 |
| *Chorthippus binotatus* | 10.41 | 12.31 | Jun-Oct | 5 | Plant | 0-2500 |
| *Chorthippus jucundus* | 12.22 | 18.05 | Jun-Oct | 5 | Plant | 200-2300 |
| *Chorthippus jacobsi* | 9.86 | 12.40 | Apr-Nov | 8 | Plant | 0-2000 |
| *Chorthippus montanus* | 8.83 | 11.20 | Jun-Sep | 4 | Plant | 1800-2300 |
| *Chorthippus nevadensis* | 7.30 | 8.53 | Ago-Sep | 2 | Plant | 1500-3300 |
| *Chorthippus parall. erythropus* | 9.75 | 11.71 | June-Nov | 6 | Plant | 100-2300 |
| *Chorthippus saulcyi moralesi* | 9.14 | 11.10 | Jul-Oct | 4 | Plant | 900-2400 |
| *Chorthippus saulcyi saulcyi* | 10.36 | 12.39 | Jul-Oct | 4 | Plant | 700-1800 |
| *Chorthippus vagans* | 8.70 | 10.58 | Apr-Nov | 8 | Plant | 0-2500 |
| *Chorthippus yersini* | 9.53 | 12.76 | May-Nov | 7 | Plant | 1200-2100 |
| *Cophopodisma pyrenaea* | 8.81 | 10.64 | Jul-Oct | 4 | Ground | 1800-2300 |
| *Dericorys carthagonovae* | 9.96 | 13.24 | Jun-Oct | 5 | Plant | 0-700 |
| *Dociostaurus crassiusculus* | 9.62 | 12.24 | Jun-Ago | 3 | Ground | 600-1000 |
| *Dociostaurus hispanicus* | 9.47 | 12.01 | Apr-Sep | 6 | Ground | 400-1500 |
| *Dociostaurus jagoi* | 7.73 | 11.02 | Jul-Nov | 5 | Ground | 0-2000 |
| *Dociostaurus maroccanus* | 13.04 | 13.46 | Jul-Ago | 2 | Ground | 400-1600 |
| *Euchorthippus chopardi* | 10.09 | 11.67 | Jun-Nov | 7 | Plant | 100-2000 |
| *Euchorthippus declivus* | 9.76 | 13.42 | Jun-Oct | 5 | Plant | 600-1400 |
| *Euchorthippus elegantulus* | 10.32 | 13.76 | Jul-Oct | 4 | Plant | 0-1800 |
| *Euthystira brachyptera* | 10.18 | 12.10 | Jun-Sept | 4 | Plant | 1300-2100 |
| *Eyprepocnemis plorans* | 16.39 | 22.83 | May-Oct | 6 | Plant | 0-100 |
| *Gomphoceridius brevipennis* | 8.87 | 10.17 | Aug-Oct | 3 | Ground | 2000-2500 |
| *Gomphocerus sibiricus* | 10.58 | 12.46 | Jul-Sep | 3 | Plant | 1800-2100 |
| *Heteracris adspersa* | 13.53 | 16.55 | Jul-Oct | 4 | Ground | 0-500 |
| *Heteracris littoralis* | 15.35 | 22.16 | Jul-Oct | 4 | Ground | 0-100 |
| *Jacobsiella imitans* | 8.91 | 11.13 | Jun-Sept | 4 | Ground | 0-100 |
| *Mioscirtus wagneri maghrebi* | 8.38 | 11.40 | Jun-Nov | 6 | Plant | 0-1500 |
| *Mioscirtus wagneri wagneri* | 7.74 | 10.86 | Jun-Nov | 6 | Plant | 0-1500 |
| *Morphacris fasciata* | 10.54 | 13.87 | Jul-Oct | 4 | Ground | 0-100 |
| *Myrmeleotettix maculatus* | 8.02 | 9.35 | Jul-Oct | 4 | Ground | 1100-2200 |
| *Oedaleus decorus* | 14.35 | 16.51 | Apr-Oct | 7 | Ground | 400-1500 |
| *Oedipoda caerulea* | 10.76 | 11.71 | Jun-Dec | 7 | Ground | 200-2000 |
| *Oedipoda charpentieri* | 9.13 | 11.72 | Jun-Dec | 7 | Ground | 100-2000 |
| *Oedipoda fuscocincta* | 11.48 | 13.56 | Jun-Oct | 5 | Ground | 600-2100 |
| *Omocestus antigai* | 8.96 | 11.25 | Jul-Oct | 4 | Ground | 1600-2100 |
| *Omocestus bolivari* | 7.23 | 8.53 | Jul-Oct | 4 | Plant | 1500-2500 |
| *Omocestus femoralis* | 9.18 | 10.86 | Jun-Oct | 5 | Plant | 1500-2000 |
| *Omocestus haemorrhoidalis* | 7.95 | 9.96 | Jul-Oct | 4 | Plant | 1500-2000 |
| *Omocestus minutissimus* | 7.53 | 7.94 | Jul-Oct | 4 | Ground | 100-2500 |
| *Omocestus navasi* | 10.22 | 12.53 | Jul-Oct | 4 | Ground | 1200-1700 |
| *Omocestus panteli* | 7.58 | 10.99 | Apr-Nov | 8 | Plant | 400-1900 |
| *Omocestus raymondi* | 8.48 | 10.51 | Jan-Dec | 12 | Ground | 0-1900 |
| *Omocestus rufipes* | 8.68 | 11.26 | Apr-Dec | 9 | Plant | 1000-2000 |
| *Omocestus uhagonii* | 7.82 | 9.37 | Jun-Oct | 5 | Ground | 1700-2200 |
| *Omocestus viridulus viridulus* | 9.03 | 11.44 | Jun-Oct | 5 | Plant | 1300-2200 |
| *Omocestus viridulus kaestneri* | 10.09 | 13.65 | Jul-Oct | 4 | Plant | 500-2100 |
| *Paracaloptenus bolivari* | 9.82 | 13.94 | Jun-Oct | 5 | Ground | 1000-1600 |
| *Paracinema tricolor* | 11.96 | 19.50 | Jul-Oct | 6 | Plant | 0-1500 |
| *Pezotettix giornae* | 6.60 | 7.63 | Jun-Nov | 6 | Ground | 0-2200 |
| *Podisma carpetana carpetana* | 10.65 | 13.01 | Jul-Sep | 3 | Ground | 1800-2000 |
| *Podisma carpetana ignatii* | 9.82 | 12.05 | Jul-Sep | 3 | Ground | 1800-2000 |
| *Podisma pedestris* | 10.38 | 12.50 | Jul-Oct | 6 | Ground | 1200-2600 |
| *Pseudosphin. azurescens* | 8.95 | 10.67 | Jun-Oct | 5 | Ground | 0-1700 |
| *Psophus stridulus* | 14.29 | 15.42 | Jul-Sep | 3 | Ground | 600-1600 |
| *Pyrgomorpha conica (\*)* | 7.20 | 10.34 | Mar-Jul | 6 | Plant | 0-1000 |
| *Ramburiella hispanica* | 11.67 | 15.49 | Jun-Oct | 5 | Plant | 0-1600 |
| *Stauroderus scalaris* | 11.39 | 13.92 | Jun-Oct | 5 | Plant | 900-2200 |
| *Stenobothrus bolivarii* | 10.62 | 14.63 | May-Ago | 4 | Plant | 700-1300 |
| *Stenobothrus festivus* | 9.72 | 10.92 | May-Ago | 4 | Plant | 0-2000 |
| *Stenobothrus grammicus* | 10.05 | 13.51 | Jun-Oct | 5 | Ground | 1100-2000 |
| *Stenobothrus lineatus* | 10.99 | 14.48 | Jun-Sep | 4 | Plant | 1300-2200 |
| *Stenobothrus nigromaculatus* | 8.97 | 10.77 | Jul-Oct | 4 | Plant | 1100-2800 |
| *Stenobothrus fischeri* | 9.75 | 12.97 | Jun-Sep | 4 | Plant | 800-1000 |
| *Stenobothrus stigmaticus* | 7.53 | 9.53 | Jul-Nov | 5 | Ground | 500-2200 |
| *Stethophyma grossum* | 12.55 | 14.01 | Jul-Sep | 3 | Ground | 1100-2000 |
| *Tropidopola cylindrica* | 10.89 | 11.06 | April-Nov | 8 | Plant | 0-500 |
| *Truxalis nasuta* | 26.70 | 45.00 | Apr-Sep | 6 | Ground | 0-700 |

(continuation)
(\*) Acridoidea

**Table S3.** Association of each grasshopper species with the nine most common habitats in which these can be found, where 0 = species not recorded in the habitat, 1 = minor association, 2 = moderate association, and 3 = strong association between the species and the habitat.

(continuation) (\*) Acridoidea
| Taxa | PDI | Habitat |
| --- | --- | --- |
| *Acrida ungarica mediterranea* | 1.000 | Wetlands, grassy river sides, marshes, reedbeds |
| *Acrotylus insubricus* | 0.642 | Steppes, dry grasslands, open habitats with disperse shrubs |
| *Acrotylus fischeri* | 1.000 | High mountain vegetation, scrublands |
| *Acrotylus patruelis* | 0.677 | Sandy soils with scarce vegetation, beaches, dunes and deserts |
| *Aiolopus puissanti* | 0.702 | Marshes, dunes, sandy wastelands |
| *Aiolopus strepens* | 0.857 | Steppes, dry grasslands, open habitats with disperse shrubs, hillsides |
| *Aiolopus thalassinus* | 0.773 | Wetlands, grassy river sides, marshes, reedbeds |
| *Arcyptera fusca* | 1.000 | Nutrient-poor grasslands, subalpine meadows, pastures, rocky slopes |
| *Arcyptera microptera* | 0.785 | Steppes and dry and warm grasslands |
| *Arcyptera tornosi* | 0.857 | Shrublands |
| *Arcyptera brevipennis* | 0.815 | Erinaceae shrubs |
| *Brachycrotaphus tryxalicerus* | 0.857 | Steppes, savanna-like habitats |
| *Bohemanella frigida* | 0.714 | Alpine meadows, stony meadows and pastures |
| *Calephorus compressicornis* | 0.487 | Wetlands, dunes, grassy river sides, marshes, reedbeds |
| *Calliptamus barbarus* | 0.481 | Steppes, dry grasslands, open habitats with disperse shrubs, stony slopes |
| *Calliptamus italicus* | 1.000 | Nutrient-poor grasslands, sand areas, rocky slopes and steppe-like places |
| *Calliptamus wattenwylanus* | 0.534 | Wastelands, rocky and grazed areas |
| *Chorthippus apicalis* | 0.857 | Grassy riversides, grassy wetlands, pastures, grasslands |
| *Chorthippus binotatus* | 0.630 | Legume shrub (scrub) lands exclusively |
| *Chorthippus jucundus* | 0.785 | Moist soil, lowland wetlands, grassy riversides, small streams (*Juncus*) |
| *Chorthippus jacobsi* | 0.773 | Steppes, dry grasslands, pastures, open habitats with disperse shrubs |
| *Chorthippus montanus* | 0.815 | Wet meadows, moors, fens, sedge reeds, bogs |
| *Chorthippus nevadensis* | 0.730 | High mountain vegetation, broom communities (piornal) |
| *Chorthippus parall. erythropus* | 0.887 | Pastures, grasslands, meadows, gardens |
| *Chorthippus saulcyi moralesi* | 0.815 | Sub-mediterranean shrub formations, mesophile montane grasslands |
| *Chorthippus saulcyi saulcyi* | 0.767 | Mesophilic grasslands |
| *Chorthippus vagans* | 0.815 | Forest ecotones, gappy grasslands, sandy pinewoods, alpine scrublands |
| *Chorthippus yersini* | 0.887 | Alpine meadows, short and dense shrublands |
| *Cophopodisma pyrenaea* | 0.857 | Alpine and subalpine meadows |
| *Dericorys carthagonovae* | 0.887 | Desert scrub, saltworts |
| *Dociostaurus crassiusculus* | 0.887 | Steppes, dry grasslands, open areas with disperse shrubs, gypsum soils |
| *Dociostaurus hispanicus* | 0.928 | Dry land pastures, xerophilic vegetation, low and sparse shrubs |
| *Dociostaurus jagoi* | 0.677 | Steppes, dry grasslands, open habitats with disperse shrubs |
| *Dociostaurus maroccanus* | 0.887 | Cereal crops, floodlands, meadows |
| *Euchorthippus chopardi* | 0.857 | Maquis, rosemary scrubs, grasslands, open habitats with disperse shrubs |
| *Euchorthippus declivus* | 0.443 | Steppes, dry grasslands, wet meadows and managed pastures |
| *Euchorthippus elegans* | 0.928 | Pastures |
| *Euthystira brachyptera* | 0.857 | Mires, damp pastures, marshy meadows, peatbogs |
| *Eyprepocnemis plorans* | 1.000 | Marshes, river streams, rice fields |
| *Gomphoceridius brevipennis* | 0.767 | Alpine meadow |
| *Gomphocerus sibiricus* | 0.815 | Pastures, alpine and subalpine meadows, rupicolous shrubs, screes |
| *Heteracris adspersa* | 0.928 | Hipersaline lagoons, dunes, beaches |
| *Heteracris littoralis* | 0.928 | Dunes, beaches, saltwort |
| *Jacobsiella imitans* | 0.928 | Movable dunes, sandy soils, pinewoods, thickets |
| *Mioscirtus wagneri maghrebi* | 1.000 | Halophilous communities, lagoon shores, low grounds with *Suaeda vera* |
| *Mioscirtus wagneri wagneri* | 0.534 | Hypersaline low ground, lagoon shores |
| *Morphacris fasciata* | 0.819 | Dunes, beaches, sandy soils |
| *Myrmeleotettix maculatus* | 0.928 | Alpine meadows, piornal, dense shrublands, stony pastures, grasslands |
| *Oedaleus decorus* | 1.000 | Rockrose scrub, heathland, meadows and cereal field margins |
| *Oedipoda caerulescens* | 0.767 | Steppes, dunes, heathlands, dry grasslands, limestone rocks |
| *Oedipoda charpentieri* | 0.857 | Rosemary scrub, thyme bushes |
| *Oedipoda fuscocincta* | 0.815 | Open habitats with disperse shrubs, broom communities, alpine meadows |
| *Omocestus antigai* | 0.815 | Thorn scrub patches, broom communities |
| *Omocestus bolivari* | 0.815 | Alpine meadows, broom communities |
| *Omocestus femoralis* | 1.000 | Alpine meadows, broom communities |
| *Omocestus haemorrhoidalis* | 0.631 | Alpine meadows, gappy and grazed pastures, pet boags, open areas |
| *Omocestus minutissimus* | 1.000 | Thorn scrub patches, broom communities |
| *Omocestus navasi* | 1.000 | Thorn scrub patches, broom communities |
| *Omocestus panteli* | 1.000 | Moist soil, lowland wetlands, grassy riversides, small streams |
| *Omocestus raymondi* | 0.730 | Open habitats with disperse shrubs, gappy grasslands, rocky slopes |
| *Omocestus rufipes* | 0.443 | Heath bogs, dry fens, old quarries, wood clearings |
| *Omocestus uhagonii* | 0.639 | Open habitats with disperse shrubs, and rocky slopes |
| *Omocestus viridulus viridulus* | 0.391 | Alpine and subalpine meadows, fens |
| *Omocestus viridulus kaestneri* | 1.000 | Humid grasslands and montane meadows |
| *Paracaloptenus bolivari* | 1.000 | Stony and rocky embankments, stony pastures |
| *Paracinema tricolor* | 0.928 | Moist soil, lowland wetlands, grassy riversides, small streams |
| *Pezotettix giornae* | 0.887 | Open habitats, shrublands, grasslands, ecotones, embankments, edges |
| *Podisma carpetana carpetana* | 1.000 | Alpine meadows, screes, rupicolous shrubs, rocky slopes |
| *Podisma carpetana ignatii* | 0.338 | Alpine meadows, screes, rupicolous shrubs, rocky slopes |
| *Podisma pedestris* | 1.000 | Alpine meadows, screes, rupicolous shrubs, rocky slopes |
| *Pseudosphin. azurescens* | 0.815 | Dry and sandy soils, gypsum soils with disperse shrub |
| *Psophus stridulus* | 0.887 | Heathland, damp pasture, subalpine meadow, clear and sunny pine forests |
| *Pyrgormorpha conica* (*) | 0.767 | Gappy habitats with open soil spots, gravel areas along rivers, dunes |
| *Ramburiella hispanica* | 1.000 | Sparto formations, halophylous communities, grasslands, dry river beds |
| *Stauroderus scalaris* | 0.887 | Warm and dry grasslands, pastures and extensive meadows, screes |
| *Stenobothrus bolivarii* | 1.000 | Dense scrublands |
| *Stenobothrus festivus* | 1.000 | Mediterranean scrubland (*Thyme* bushes) |
| *Stenobothrus grammicus* | 0.677 | Alpine meadows, pastures, thicket, rosemary scrub |
| *Stenobothrus lineatus* | 0.857 | Heathland, grassy margins of damp woodland, mountain meadows |
| *Stenobothrus nigromaculatus* | 0.631 | Grappy and grazed grasslands, subalpine meadows |
| *Stenobothrus fischeri* | 1.000 | Gappy, dry and poor-nutrient soils |
| *Stenobothrus stigmaticus* | 0.702 | Heathland, grazed grasslands, alpine meadows, dense shrublands |
| *Stethophyma grossum* | 1.000 | Alpine meadows, pastures near rivers, pet boags |
| *Tropidopola cylindrica* | 0.857 | Reedbed, river streams, saltworts |
| *Truxalis nasuta* | 0.624 | Gappy scrublands, sandy soils |

(continuation)
(\*) Acridoidea

**Table S4.**
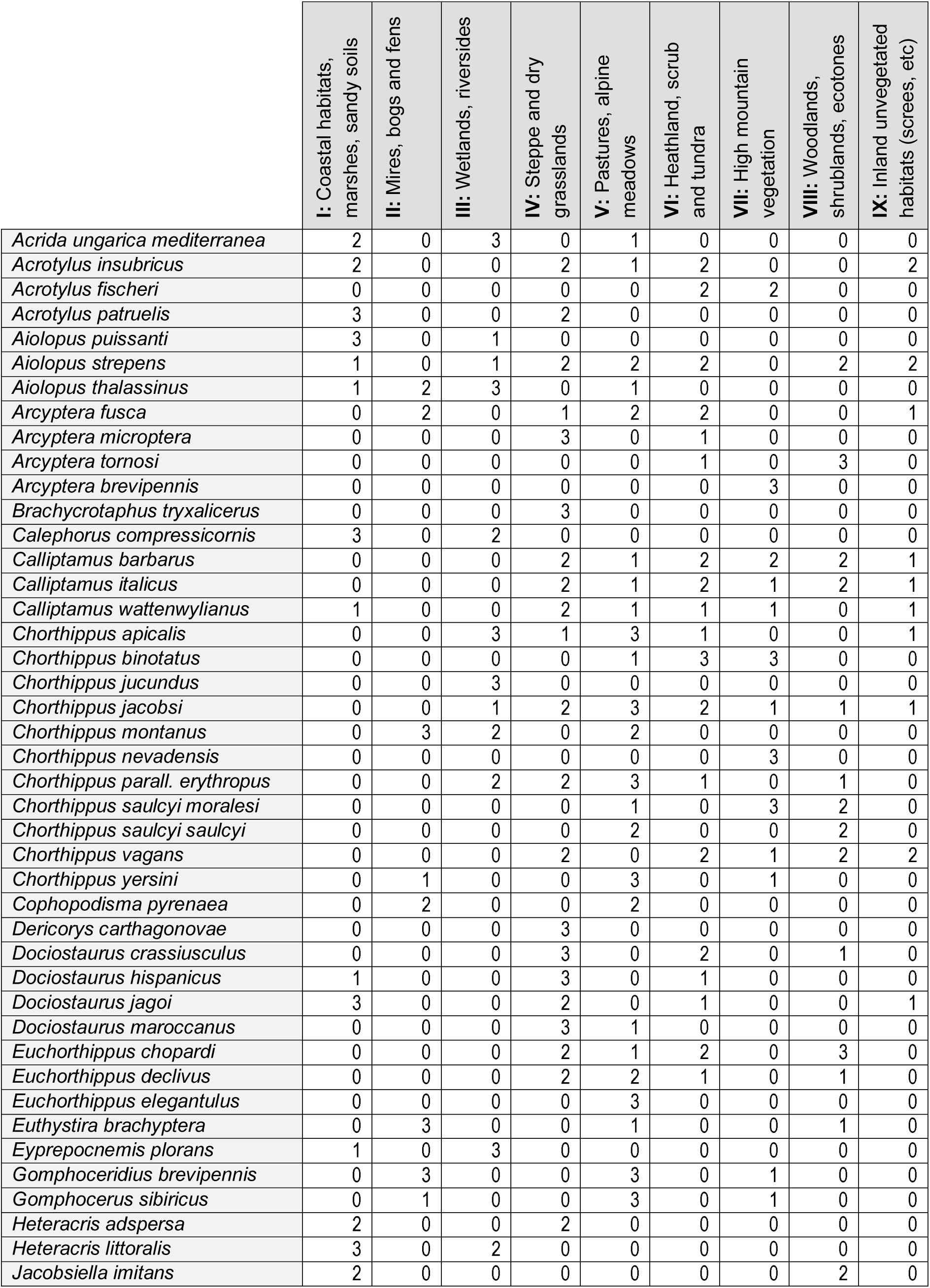
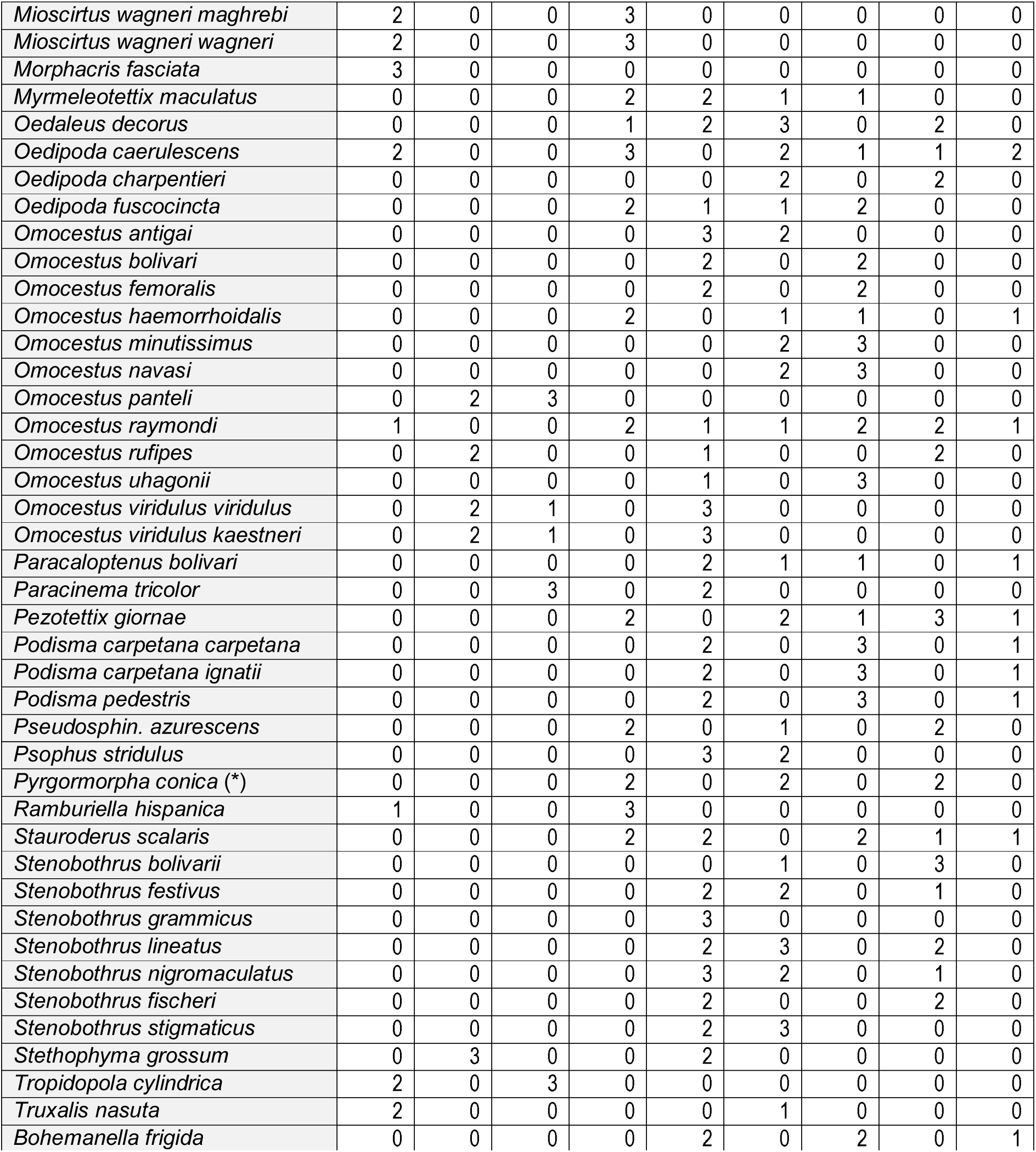

(continuation) (\*) Acridoidea
|  | I: Coastal habitats,<br>marshes, sandy soils | II: Mires, bogs and fens | III: Wetlands, riversides | IV: Steppe and dry<br>grasslands | V: Pastures, alpine<br>meadows | VI: Heathland, scrub<br>and tundra | VII: High mountain<br>vegetation | VIII: Woodlands,<br>shrublands, ecotones | IX: Inland unvegetated<br>habitats (scree, etc) |
| --- | --- | --- | --- | --- | --- | --- | --- | --- | --- |
| *Acrida ungarica mediterranea* | 2 | 0 | 3 | 0 | 1 | 0 | 0 | 0 | 0 |
| *Acrotylus insubricus* | 2 | 0 | 0 | 2 | 1 | 2 | 0 | 0 | 2 |
| *Acrotylus fischeri* | 0 | 0 | 0 | 0 | 0 | 2 | 2 | 0 | 0 |
| *Acrotylus patruelis* | 3 | 0 | 0 | 2 | 0 | 0 | 0 | 0 | 0 |
| *Aiolopus puissantii* | 3 | 0 | 1 | 0 | 0 | 0 | 0 | 0 | 0 |
| *Aiolopus strepens* | 1 | 0 | 1 | 2 | 2 | 2 | 0 | 2 | 2 |
| *Aiolopus thalassinus* | 1 | 2 | 3 | 0 | 1 | 0 | 0 | 0 | 0 |
| *Arcyptera fusca* | 0 | 2 | 0 | 1 | 2 | 2 | 0 | 0 | 1 |
| *Arcyptera microptera* | 0 | 0 | 0 | 3 | 0 | 1 | 0 | 0 | 0 |
| *Arcyptera tornosi* | 0 | 0 | 0 | 0 | 0 | 1 | 0 | 3 | 0 |
| *Arcyptera brevipennis* | 0 | 0 | 0 | 0 | 0 | 0 | 3 | 0 | 0 |
| *Brachycrotaphus tryxalicerus* | 0 | 0 | 0 | 3 | 0 | 0 | 0 | 0 | 0 |
| *Calephorus compressicornis* | 3 | 0 | 2 | 0 | 0 | 0 | 0 | 0 | 0 |
| *Calliptamus barbarus* | 0 | 0 | 0 | 2 | 1 | 2 | 2 | 2 | 1 |
| *Calliptamus italicus* | 0 | 0 | 0 | 2 | 1 | 2 | 1 | 2 | 1 |
| *Calliptamus wattenwylanus* | 1 | 0 | 0 | 2 | 1 | 1 | 1 | 0 | 1 |
| *Chorthippus apicalis* | 0 | 0 | 3 | 1 | 3 | 1 | 0 | 0 | 1 |
| *Chorthippus binotatus* | 0 | 0 | 0 | 0 | 1 | 3 | 3 | 0 | 0 |
| *Chorthippus jucundus* | 0 | 0 | 3 | 0 | 0 | 0 | 0 | 0 | 0 |
| *Chorthippus jacobsi* | 0 | 0 | 1 | 2 | 3 | 2 | 1 | 1 | 1 |
| *Chorthippus montanus* | 0 | 3 | 2 | 0 | 2 | 0 | 0 | 0 | 0 |
| *Chorthippus nevadensis* | 0 | 0 | 0 | 0 | 0 | 0 | 3 | 0 | 0 |
| *Chorthippus parall. erythropus* | 0 | 0 | 2 | 2 | 3 | 1 | 0 | 1 | 0 |
| *Chorthippus saulcyi moralesi* | 0 | 0 | 0 | 0 | 1 | 0 | 3 | 2 | 0 |
| *Chorthippus saulcyi saulcyi* | 0 | 0 | 0 | 0 | 2 | 0 | 0 | 2 | 0 |
| *Chorthippus vagans* | 0 | 0 | 0 | 2 | 0 | 2 | 1 | 2 | 2 |
| *Chorthippus yersini* | 0 | 1 | 0 | 0 | 3 | 0 | 1 | 0 | 0 |
| *Cophopodisma pyrenaica* | 0 | 2 | 0 | 0 | 2 | 0 | 0 | 0 | 0 |
| *Dericorys carthagonovae* | 0 | 0 | 0 | 3 | 0 | 0 | 0 | 0 | 0 |
| *Dociostaurus crassiusculus* | 0 | 0 | 0 | 3 | 0 | 2 | 0 | 1 | 0 |
| *Dociostaurus hispanicus* | 1 | 0 | 0 | 3 | 0 | 1 | 0 | 0 | 0 |
| *Dociostaurus jagoi* | 3 | 0 | 0 | 2 | 0 | 1 | 0 | 0 | 1 |
| *Dociostaurus maroccanus* | 0 | 0 | 0 | 3 | 1 | 0 | 0 | 0 | 0 |
| *Euchorthippus chopardi* | 0 | 0 | 0 | 2 | 1 | 2 | 0 | 3 | 0 |
| *Euchorthippus declivus* | 0 | 0 | 0 | 2 | 2 | 1 | 0 | 1 | 0 |
| *Euchorthippus elegantulus* | 0 | 0 | 0 | 0 | 3 | 0 | 0 | 0 | 0 |
| *Euthystira brachyptera* | 0 | 3 | 0 | 0 | 1 | 0 | 0 | 1 | 0 |
| *Eyprepocnemis plorans* | 1 | 0 | 3 | 0 | 0 | 0 | 0 | 0 | 0 |
| *Gomphoceridius brevipennis* | 0 | 3 | 0 | 0 | 3 | 0 | 1 | 0 | 0 |
| *Gomphocerus sibiricus* | 0 | 1 | 0 | 0 | 3 | 0 | 1 | 0 | 0 |
| *Heteracris adspersa* | 2 | 0 | 0 | 2 | 0 | 0 | 0 | 0 | 0 |
| *Heteracris littoralis* | 3 | 0 | 2 | 0 | 0 | 0 | 0 | 0 | 0 |
| *Jacobsiella imitans* | 2 | 0 | 0 | 0 | 0 | 0 | 0 | 2 | 0 |
| *Mioscirtus wagneri maghrebi* | 2 | 0 | 0 | 3 | 0 | 0 | 0 | 0 | 0 |
| *Mioscirtus wagneri wagneri* | 2 | 0 | 0 | 3 | 0 | 0 | 0 | 0 | 0 |
| *Morphacris fasciata* | 3 | 0 | 0 | 0 | 0 | 0 | 0 | 0 | 0 |
| *Myrmeleotettix maculatus* | 0 | 0 | 0 | 2 | 2 | 1 | 1 | 0 | 0 |
| *Oedaleus decorus* | 0 | 0 | 0 | 1 | 2 | 3 | 0 | 2 | 0 |
| *Oedipoda caerulescens* | 2 | 0 | 0 | 3 | 0 | 2 | 1 | 1 | 2 |
| *Oedipoda charpentieri* | 0 | 0 | 0 | 0 | 0 | 2 | 0 | 2 | 0 |
| *Oedipoda fuscocincta* | 0 | 0 | 0 | 2 | 1 | 1 | 2 | 0 | 0 |
| *Omocestus antigai* | 0 | 0 | 0 | 0 | 3 | 2 | 0 | 0 | 0 |
| *Omocestus bolivari* | 0 | 0 | 0 | 0 | 2 | 0 | 2 | 0 | 0 |
| *Omocestus femoralis* | 0 | 0 | 0 | 0 | 2 | 0 | 2 | 0 | 0 |
| *Omocestus haemorrhoidalis* | 0 | 0 | 0 | 2 | 0 | 1 | 1 | 0 | 1 |
| *Omocestus minutissimus* | 0 | 0 | 0 | 0 | 0 | 2 | 3 | 0 | 0 |
| *Omocestus navasi* | 0 | 0 | 0 | 0 | 0 | 2 | 3 | 0 | 0 |
| *Omocestus panteli* | 0 | 2 | 3 | 0 | 0 | 0 | 0 | 0 | 0 |
| *Omocestus raymondi* | 1 | 0 | 0 | 2 | 1 | 1 | 2 | 2 | 1 |
| *Omocestus rufipes* | 0 | 2 | 0 | 0 | 1 | 0 | 0 | 2 | 0 |
| *Omocestus uhagonii* | 0 | 0 | 0 | 0 | 1 | 0 | 3 | 0 | 0 |
| *Omocestus viridulus viridulus* | 0 | 2 | 1 | 0 | 3 | 0 | 0 | 0 | 0 |
| *Omocestus viridulus kaestneri* | 0 | 2 | 1 | 0 | 3 | 0 | 0 | 0 | 0 |
| *Paracaloptenus bolivari* | 0 | 0 | 0 | 0 | 2 | 1 | 1 | 0 | 1 |
| *Paracinema tricolor* | 0 | 0 | 3 | 0 | 2 | 0 | 0 | 0 | 0 |
| *Pezotettix giornae* | 0 | 0 | 0 | 2 | 0 | 2 | 1 | 3 | 1 |
| *Podisma carpetana carpetana* | 0 | 0 | 0 | 0 | 2 | 0 | 3 | 0 | 1 |
| *Podisma carpetana ignatii* | 0 | 0 | 0 | 0 | 2 | 0 | 3 | 0 | 1 |
| *Podisma pedestris* | 0 | 0 | 0 | 0 | 2 | 0 | 3 | 0 | 1 |
| *Pseudosphing. azurescens* | 0 | 0 | 0 | 2 | 0 | 1 | 0 | 2 | 0 |
| *Psophus stridulus* | 0 | 0 | 0 | 0 | 3 | 2 | 0 | 0 | 0 |
| *Pyrgormorpha conica (\*)* | 0 | 0 | 0 | 2 | 0 | 2 | 0 | 2 | 0 |
| *Ramburiella hispanica* | 1 | 0 | 0 | 3 | 0 | 0 | 0 | 0 | 0 |
| *Stauroderus scalaris* | 0 | 0 | 0 | 2 | 2 | 0 | 2 | 1 | 1 |
| *Stenobothrus bolivarii* | 0 | 0 | 0 | 0 | 0 | 1 | 0 | 3 | 0 |
| *Stenobothrus festivus* | 0 | 0 | 0 | 0 | 2 | 2 | 0 | 1 | 0 |
| *Stenobothrus grammicus* | 0 | 0 | 0 | 0 | 3 | 0 | 0 | 0 | 0 |
| *Stenobothrus lineatus* | 0 | 0 | 0 | 0 | 2 | 3 | 0 | 2 | 0 |
| *Stenobothrus nigromaculatus* | 0 | 0 | 0 | 0 | 3 | 2 | 0 | 1 | 0 |
| *Stenobothrus fischeri* | 0 | 0 | 0 | 0 | 2 | 0 | 0 | 2 | 0 |
| *Stenobothrus stigmaticus* | 0 | 0 | 0 | 0 | 2 | 3 | 0 | 0 | 0 |
| *Stethophyma grossum* | 0 | 3 | 0 | 0 | 2 | 0 | 0 | 0 | 0 |
| *Tropidopola cylindrica* | 2 | 0 | 3 | 0 | 0 | 0 | 0 | 0 | 0 |
| *Truxalis nasuta* | 2 | 0 | 0 | 0 | 0 | 1 | 0 | 0 | 0 |
| *Bohemanella frigida* | 0 | 0 | 0 | 0 | 2 | 0 | 2 | 0 | 1 |

**Fig. S1.**
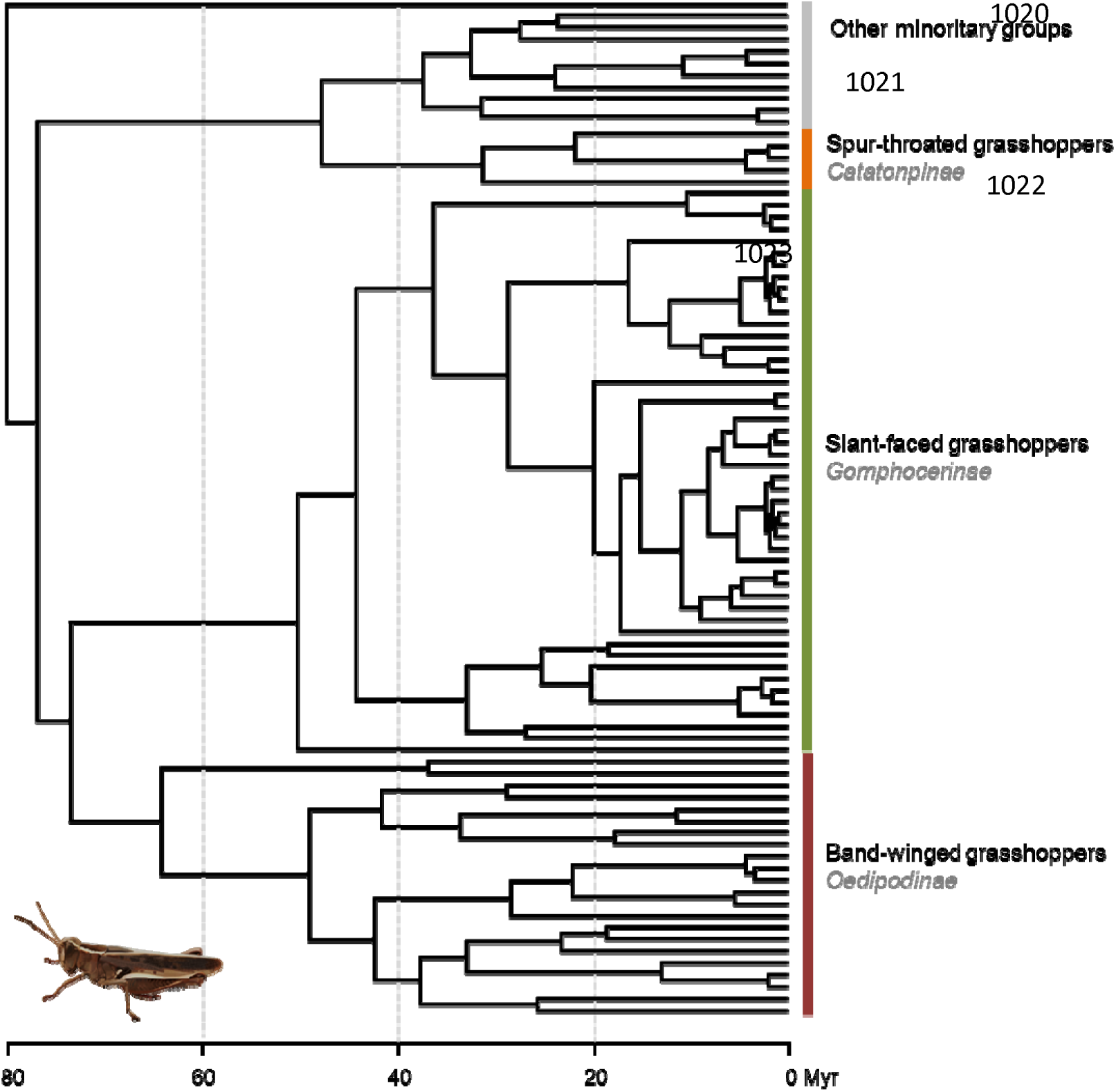
The Bayesian maximum clade credibility (MCC) tree.

